# LOOP-TAG: massively-parallel measurement of length-dependent protein-mediated DNA looping probabilities by tagmentation *in vitro*

**DOI:** 10.64898/2026.09.21.753159

**Authors:** Naomi R. Kennel, Nicole A. Becker, Justin P. Peters, L. James Maher

## Abstract

We are interested in DNA stiffness *in vitro* and *in vivo*. Classical approaches for measuring this parameter *in vitro* involve tedious T4 DNA ligase-mediated cyclization kinetics experiments performed for one DNA fragment length at a time to establish effective local end-end concentrations (*J*-factors). Protein-mediated DNA looping is a more relevant biological reaction. An efficient approach to determine length-dependent protein-mediated DNA *J*-*loop* values *in vitro* has been lacking. We describe LOOP-TAG, a massively-parallel method. We leverage intramolecular DNA looping and tagmentation catalyzed by Tn5 transposase (Tnp) tethered to the terminus of a ∼1,000-bp bead-bound duplex DNA. We detected loops terminating with Tnp. Counts of the resulting DNA looping-dependent bead-bound tagmentation products from deep sequencing determine DNA loop length probabilities at single-base pair resolution for all bead-bound fragments. When normalized to DNA fragment length probabilities from tagmentation using known concentrations of free Tnp, length-dependent *J*-*loop* values are directly determined. We demonstrate LOOP-TAG for Tnp tethered in two ways and in the presence of DNA-kinking architectural DNA binding protein Nhp6A. Intermediate length *J-loop* values are consistent with expectations of wormlike chain theory after accounting for strong sequence-dependence of Tnp. DNA loops are shifted to smaller sizes in the presence of Nhp6A, as predicted. (200/200 words)

## Introduction

Double-helical DNA is among the stiffest biopolymers in terms of resistance to bending and twisting. Its persistence length in low-salt buffers is ∼150 bp, describing the DNA length over which the average deviation from its initial trajectory is one radian (1). Resistance to DNA twisting (twist constant ∼2 × 10^−19^ erg-cm) gives rise to a similar twist persistence length (1). We and others have used theoretical and experimental approaches in attempts to parse the relative contributions of compaction by base-pair stacking (2) vs. stretching by electrostatic repulsion (1,3) to these parameters. Counterion condensation theory (4,5) proposes that ∼76% of duplex DNA charges are neutralized by condensation of monovalent cations, and this degree of neutralization is essentially independent of bulk monovalent cation concentration *in vitro*, with the persistence length of the resulting polymer being cation concentration-dependent due to its residual anionic character (4). Despite many studies, a clear understanding of the relative contributions of electrostatics and base pair stacking to DNA stiffness remains elusive: it appears that both base pair stacking forces and electrostatic stretching forces make contributions.

The ability to measure short-range double-helical DNA stiffness is important because stiffness must be overcome for DNA compaction and looping over short distances involved in local gene regulation. DNA binding and deformation by proteins such as histone octamers (6,7) and architectural proteins (8) is substantially driven by the favorable entropy of condensed counterion release from double-helical DNA upon cationic protein binding, together with asymmetric phosphodiester charge neutralization (9–12). DNA stiffness has been measured *in vitro* and *in vivo* by both ensemble and single-molecule approaches (1,13–16). Of particular interest to our laboratory has been the comparison of DNA stiffness measurements *in vitro* and *in vivo*. DNA looping over hundreds of base pairs should be dramatically constrained by local DNA stiffness to both bending and twisting. Such effects become less significant for DNA looping at the megabase pair scale involved in genome organization (1,17,18).

*In vitro* DNA stiffness assays have classically exploited bacteriophage T4 DNA ligase-mediated cyclization kinetics experiments (Fig. 1AB). For any given DNA fragment bearing modestly cohesive termini, attention to key experimental parameters and assumptions about T4 DNA ligase concentration allows conditions to be identified where the ratio of the rates of unimolecular cyclization (Fig. 1A) to bimolecular dimerization (Fig. 1B) defines the *J*-factor, an estimate of the effective local concentration of one DNA terminus in the vicinity of the other (1). *In vitro J*-factor estimates predict that the most probable length for double-helical DNA cyclization is ∼450 bp, with cyclization probability dropping drastically for shorter segments due to DNA stiffness, and dropping gradually for longer segments due to dilution of termini (1,15). Other ensemble approaches to *in vitro* double-helical DNA stiffness quantitation have been reported (19). It should be emphasized that T4 DNA ligase cyclization assays require radioactive DNA labeling, precise DNA quantitation, attention to ligase concentration, and data fitting to rate equations. Importantly, the method can only be applied for DNA fragments above ∼110 bp where cyclization rate is detectable (14). Increasing DNA ligase to enhance cyclization yield risks erroneous conclusions about hyperflexibility of short DNA (13,14). Inability to define *J*-factors for short DNA is further limited by the observation that strained minicircles in the range of 85 bp begin to show kinking behavior (20). Thus, despite precision, T4 DNA ligase cyclization kinetics experiments are challenging, expensive and inapplicable to study DNA lengths below ∼110 bp (15). Elegant single-molecule fluorescence resonance energy transfer (FRET)-based cyclization approaches with tethered DNA fragments have been applied to study the sequence-dependence of double-helical DNA stiffness (16,21–23), though direct translation to *J*-factors can be complicated by experimental conditions including long cohesive single-stranded DNA termini.

**Fig. 1.**
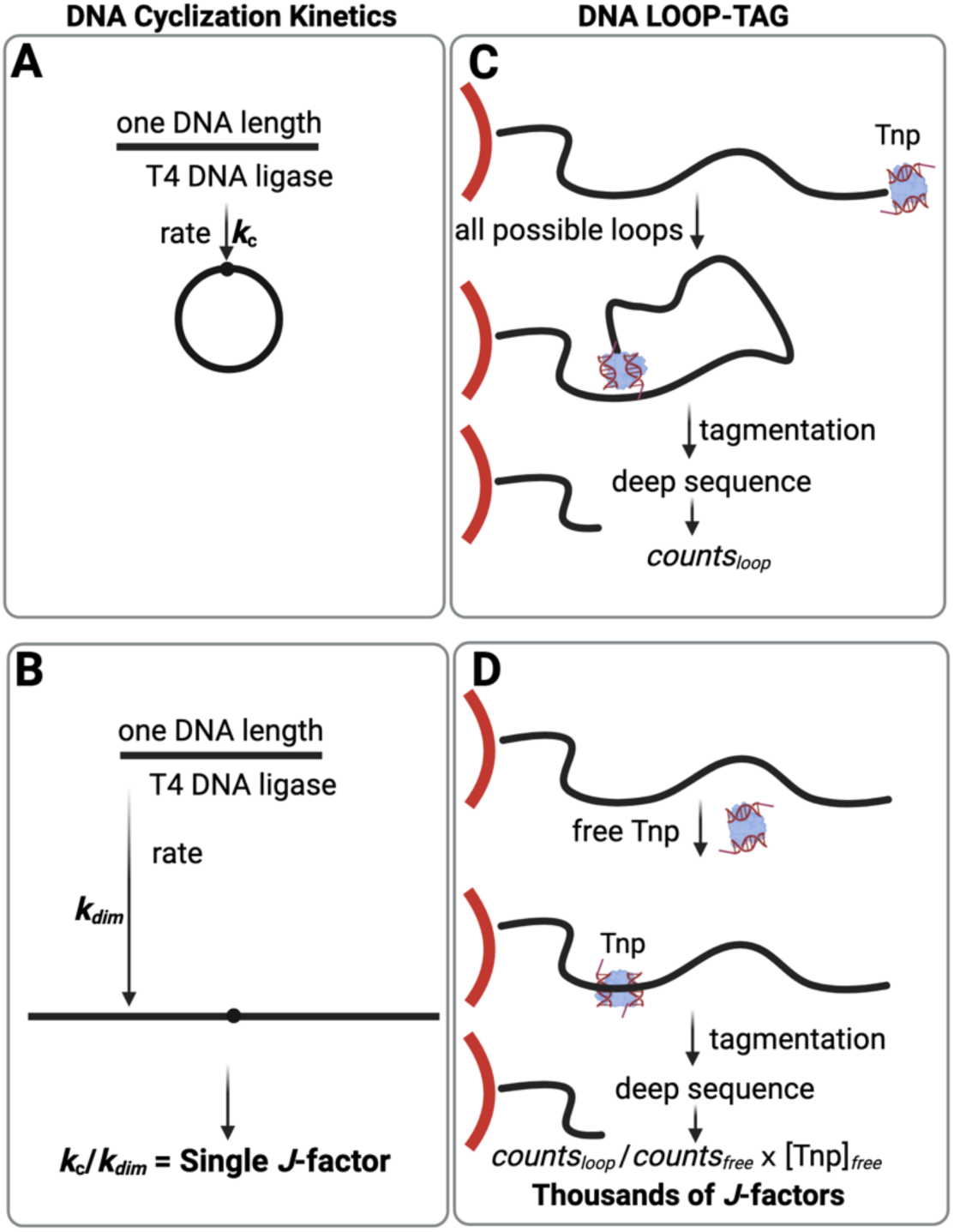
Experimental methods for estimating *J*-factors *in vitro*. A-B. Conventional T4 DNA ligase-mediated DNA cyclization kinetics. A single *J*-factor (in molar units reflecting effective DNA end-end concentration) is determined by calculating the ratio between the measured DNA concentration-independent cyclization rate (A) and the DNA concentration-dependent dimerization rate (B) for ligation of a DNA fragment of one length. C-D. Generic depiction of LOOP-TAG concept where Tnp conjugation details are not shown. Thousands of *J*-factors are simultaneously determined by protein-mediated Tn5 transposase (Tnp)-mediated intramolecular tagmentation of an immobilized DNA construct. Deep sequencing of tethered fragments is used to calculate the ratios of length-dependent counts for immobilized DNA with terminally-conjugated Tnp (C) vs. immobilized DNA exposed to free Tnp (D). Ratios are multiplied by the known free Tnp concentration to give length-dependent *J*-*loop* values.

Motivated by the work of Record et al. (24), and Müller-Hill et al. (25,26) we have also studied protein-mediated DNA looping required for maximal repression of the *E. coli lactose* operon as an approach to estimate *J*-factors *in vivo* (*J*-looping) (15). Such *in vivo J*-looping probabilities in *E. coli* are many orders of magnitude higher than *in vitro*, presumably reflecting the “softening” effects of negative DNA supercoiling, and protein bridging that reduce necessary DNA deformation, and the presence of a plethora of sequence-specific and -nonspecific architectural DNA bending and kinking proteins *in vivo* (27–29).

The present work focuses on overcoming the tedious nature of conventional T4 DNA ligase-mediated cyclization assays involving quantitative electrophoretic gel analysis of products of kinetic experiments, or single-molecule experiments involving cyclization rate measurements. These methods share a low throughput limitation in that measurements are made for one double-helical DNA fragment length at a time. We describe an alternative method that allows *J*-looping estimates *for thousands of DNA fragment lengths simultaneously*.

Further, we reasoned that measuring double-helical DNA stiffness *in vitro* by protein-mediated DNA looping is more biologically relevant than ligase-catalyzed cyclization, the latter requiring both 360° of DNA bending and helical phasing and alignment of DNA termini. We now describe a massively parallel *in vitro* protein-mediated DNA looping method, DNA LOOPing measured by TAGmentation (“LOOP-TAG”), that generates *J*-looping measurements for thousands of DNA fragment lengths *simultaneously*, exploiting the remarkable tagmentation properties of a hyperactive mutant form of *E. coli* Tn5 transposase (Tnp) (30–33), i.e. the ability of Tnp to cleave DNA and simultaneously add short adapter sequences to the cleaved termini. This reaction requires Mg^2+^, so it can be controlled by metal chelation. These Tnp properties have previously been exploited in Assay for Transposase-Accessible Chromatin using sequencing [“ATAC-seq,” (34)] and Cleavage Under Targets and Tagmentation [“CUT&Tag,” (35)] in genome-wide DNA accessibility and protein mapping studies.

## Materials and Methods

### Buffers

Experimental buffers are described in Supplemental Table S1.

### LOOP-TAG

#### DNA target synthesis

DNA sequences are provided in Supplemental Table S2. DNA primers are listed in Supplemental Tables S3-S7. DNA Target 1 (1110 bp) was created by PCR amplification from plasmid pJ1506. DNA Target 2 (1161 bp) was created by PCR amplification from plasmid pJ3054. DNA Target 3 (1033 bp) was created by PCR amplification from plasmid pJ2656. All synthetic oligonucleotides were from IDT (Coralville, IA). PCR (100 µL) included 10 ng plasmid template, 0.4 mM each forward and reverse primers, 100 µg/mL bovine serum albumin (BSA), 1× PCR buffer (Invitrogen #10342178), 4 mM MgCl_2_, 0.2 mM each dNTP, and 5 U *Taq* DNA polymerase (Invitrogen #10342178). Cycle temperatures and times were 94°C (3 min), 25 cycles of [98°C (10 s), 60°C (20 s), and 72°C (60 s)] followed by 72°C (5 min). After PCR, reactions were incubated at 37°C for 60 min with 40 U DpnI restriction enzyme (New England Biolabs #R0176S). PCR products were purified using a QIAquick PCR Purification Kit (QIAGEN #28104) and quantified using a NanoDrop 1000 Spectrophotometer (ThermoFisher).

#### Immobilizing biotinylated PCR targets

M-270 Dynabeads™ streptavidin magnetic bead suspension (Invitrogen #65-601; 100 µL) was washed twice with 500 µL 2× B&W buffer and resuspended in 1 mL B&W buffer. Terminally-biotinylated DNA target (3 ng) was added to the resuspended beads and incubated with rotation at room temperature for 30 min. Loaded M-270 DNA beads were washed three times with 500 µL B&W buffer with gentle mixing between each wash and resuspended in 400 µL Dig 300 buffer. DNA-loaded beads were stored at 4°C until use. For all subsequent annealing, antibody, and free Tnp experiments, 50 µL of the 400 µL of DNA-loaded beads was used, corresponding to 12.5 µL of the original M270 beads and 0.375 ng of DNA.

#### Adapters for Tnp conjugation by terminal Tnp target annealing

Adapter sequences are shown in Supplemental Table S6. 45 µM of both 7065/7066 (Adapter A) and 7065/7901 (Adapter B) duplexes were created by combining 22.5 µL of each 100 µM oligonucleotide stock with 5 µL 10× annealing buffer. Annealing mixtures were heated in a thermocycler to 95°C for three min and cooled to 4°C at a rate of 1°C/min.

#### Adapter loading into Tn5 Tnp (for free Tnp or terminal Tnp target annealing studies)

For experiments with free Tnp or Tnp conjugation by terminal target annealing, 5.1 µL of 75.5 µM unloaded Tagmentase (Diagenode # C01070010-20) was mixed with 13 µL 45 µM 7065/7066 mosaic adapter A duplex (for free Tnp studies) or 7065/7901 mosaic adapter A duplex (for terminal target annealing), and 30.9 µL 2× Tagmentation Buffer (Diagenode # C01019043-100) and then incubated for 1 h at room temperature. The resulting 7.7 µM solution of loaded Tnp was used promptly.

#### Adapter loading into pA-Tnp (for free Tnp or terminal Tnp target annealing studies)

For experiments with free pA-Tnp or complex of antibody-bound pA-Tnp, 7 µL of 54.9 µM pA-Tn5 Tnp - unloaded (Diagenode #C01070002-30) was mixed with 13 µL 45 µM 7065/7066 mosaic adapter A duplex and 29 µL 2× Tagmentation Buffer (Diagenode # C01019043-100) and then incubated for 1 h at room temperature. The resulting 7.7 µM solution of loaded pA-Tnp was used promptly.

#### Validation of adapter loading into Tnp protein

Freshly prepared mosaic A adapter, 7065/7066, loaded pA-Tnp and Tnp were analyzed by denaturing and native polyacrylamide gel electrophoresis to determine the efficiency of adapter A loading into the Tnp protein. pA-Tnp or Tnp were loaded with increasing molar ratios of adapter A (7065/7066) : Tnp (0:1, 0.5:1, 1.4:1, and 2:1). Identical 20-pmol pA-Tnp or Tnp samples were loaded onto one of three polyacrylamide gels. First, a 6% 29:1 acrylamide:bisacrylamide native polyacrylamide gel in 0.5× TBE (45 mM Tris-borate and 1 mM EDTA) was electrophoresed in 0.5× TBE at 4° C for 140 V for 70 min. Next, a NativePAGE™ bis-Tris 3 to 12% polyacrylamide gradient gel (ThermoFisher # BN1001box) was electrophoresed using NativePAGE™ Running Buffer Kit (1× anode buffer and 1× light blue cathode buffer; ThermoFisher # BN2007) at 4° C for 140 V for 70 min. Finally, a 10% bis-Tris SDS denaturing polyacrylamide gel (ThermoFisher # NP0301BOX) was electrophoresed in 1× MES buffer (ThermoFisher # NP0002) at room temperature for 140 V for 70 min). Gels were post-stained first with SYBR Gold for nucleic acid staining, followed by Coomassie protein stain. Gels were fixed (40% methanol, 10% acetic acid) prior to staining with 1× Sybr Gold nucleic acid stain (ThermoFisher # S11494) and imaged with a Typhoon imager (Amersham) to determine nucleic acid signal. The gels were then immediately stained with Coomassie Brilliant Blue R-250 Staining Solution (ThermoFisher # 1610436), followed by destaining according to manufacturer and imaged with a Typhoon imager (Amersham) to determine protein signal.

#### Tnp-DNA conjugation by terminal annealing

Tnp terminal conjugation to target duplexes was by terminal target single strand annealing to the complementary single-stranded sequence of the 7065/7091 mosaic A adapter. Target DNA with an appropriate 10-nt terminal single-stranded complement to the mosaic A terminal iSp9 sequence was prepared by PCR using non-extendable distal primer 7906 (Target 1) or 7907 (Target 2). Target DNA loaded (50 µL) M-270 magnetic beads were bound to a magnet and resuspended in 1 mL Dig 600 buffer containing 100 µg/mL BSA (New England Biolabs #B9200S) and 100 µg/mL salmon sperm DNA (Invitrogen #15-632-011) to suppress nonspecific Tnp binding to the body of the target DNA. An aliquot of 7.7 µM Tnp loaded with 7065/7901 mosaic A adapter duplex was diluted to 1 µM in Tagmentation Buffer 2× (Diagenode # C01019043-100). Terminal annealing involved addition of 2 µL of 1 µM loaded Tnp (2 nM final concentration) to each tube in 1 mL bead solution including BSA (100 µg/mL) and salmon sperm DNA (100 µg/mL) competitors with incubation in a water bath at 37°C for 10 min, cooling to room temperature for 10 min, and placement on ice for 10 min. To further deplete Tnp nonspecifically bound to the target DNA body rather than the terminus, DNA-beads were bound to a magnet and washed twice at room temperature with 500 µL Dig 2000 Buffer (one minute with rotation for each wash). DNA-beads were then washed with 500 µL Looping Buffer, resuspended in 1500 µL Looping Buffer and gently mixed to ensure bead dispersion.

#### Protein-A Tnp-DNA conjugation via antibody

Terminally-biotinylated target DNA was prepared by PCR to produce a distal digoxigenin-conjugated terminus. Target DNA loaded (50 µL) M-270 magnetic beads were bound to a magnet and then resuspended in an equivalent volume of ice-cold Antibody buffer containing 1 µL 0.5 mg/mL anti-digoxigenin recombinant rabbit monoclonal antibody (Invitrogen #9H27L19) per 75 µL resuspension. Bead-DNA solution was incubated at 4°C for at least 2 h, with rotation. Bead-DNA was bound to a magnet and washed twice with 1 mL Dig 300 buffer. Beads were resuspended in the original volume of Dig 300 buffer and divided into 50 µL aliquots for subsequent binding steps. 50 µL antibody-bound bead aliquots were washed once with 500 µL Dig 600 buffer and resuspended in 300 µL Dig 600 buffer containing 5 nM Adapter A (7065/7066)-loaded Protein A (pA)-Tnp and 100 µg/mL salmon sperm DNA (Invitrogen #15-632-011). Samples were incubated with rotation at room temperature for 1 h and then bound to a magnet and washed twice with 1 mL Dig 2000 (1 min wash with rotation each time). Resulting antibody-mediated Tnp-conjugated DNA beads were then washed with 500 µL T4 ligase buffer, resuspended in 1500 µL Looping Buffer and gently mixed to ensure bead dispersion.

#### Experiments involving Nhp6A

For experiments testing effects of purified recombinant yeast Nhp6A protein (15), immediately following DNA-bead resuspension in Looping Buffer, Nhp6A (10 nM final concentration diluted in Tnp Dilution Buffer with 377 µg/mL BSA) was added to LOOP-TAG reactions and allowed to incubate for 20 min before Tnp activation.

#### Tagmentation

All LOOP-TAG experiments were performed in three independent replicates with all replicates or derived statistics shown, as indicated. Tnp activation was with a MgCl_2_ concentration 1 mM higher than any EDTA in the buffer. Solutions of terminally-biotinylated target DNAs immobilized on streptavidin beads in Looping Buffer were activated with MgCl_2_ for 10 min (antibody-conjugated Tnp and annealed Tnp) with rotation at room temperature. Tagmentation reactions were stopped by addition of 0.1% SDS and 10 mM EDTA. To ensure complete reaction inactivation, samples were placed on a heating block at 70°C for 20 min. DNA-beads were washed twice with 500 µL TE buffer supplemented with 0.1% Triton-X Buffer and resuspended in 12.5 µL H_2_O.

#### Post-tagmentation PCR

Samples of resuspended bead-DNA (6 µL) were used in 25 µL PCR reactions for validation controls and analyzed by gel electrophoresis. Samples of resuspended bead-DNA (12 µL) were used in 50 µL PCR amplifications prior to sequencing. Reactions included one-half volume FailSafe™ PCR 2× PreMix I (Lucigen # FSP995I), 0.4 mM forward and reverse primers, and 2.5 U FailSafe™ Enzyme Mix (Lucigen # FSE51100). Cycle conditions were 94°C (3 min), 22 cycles of [94°C (30 s), 63°C (30 s), and 72°C (60 s)], followed by 72°C (5 min). Unlike conventional Tnp tagmentation reactions where both DNA fragment termini arise from Tnp adapter insertions requiring an initial DNA polymerase extension step for repair of termini to enable PCR with adapter primers, the LOOP-TAG protocol benefits from a defined anchor primer binding site at the bead terminus. Post-tagmentation PCR thus initiates with a denaturation step and the distal mosaic end sequence is polymerized into a full blunt-ended PCR-compatible duplex after denaturation, annealing of the bead-terminal primer, and extension of the bottom strand.

#### Confirmation of minimal nonspecific Tnp binding to target DNA body

Multiple control samples were included in all LOOP-TAG protocols to ensure minimal nonspecific Tnp body binding during LOOP-TAG reactions. Each control excluded one individual LOOP-TAG component including terminally-biotinylated target DNAs immobilized on M-270 streptavidin magnetic beads, anti-Dig antibody, pA-Tnp, Tnp, or MgCl_2_ activation. A control to eliminate annealed Tnp conjugates involved denaturation at 37°C for 10 min, then rapid cooling on ice. DNA targets lacking a terminal single strand (preventing annealed Tnp conjugation) or lacking terminal digoxygenin (preventing antibody-mediated pA-Tnp conjugation) determined background signal for the corresponding LOOP-TAG reactions. All validation controls were subjected to electrophoresis through a 2% agarose gel and imaged with a Typhoon imager (Amersham) to determine PCR amplification signal.

#### Characterization and optimization of free Tnp and free pA-Tnp tagmentation

50 µL of the DNA-bead preparations used in LOOP-TAG reactions were resuspended in 1500 µL Looping Buffer. Tnp (2 nM, 0.4 nM, or 0.08 nM diluted in Tagmentation Buffer 2× (Diagenode # C01019043-100) containing 100 µg/mL BSA) were added to each sample in triplicate and allowed to bind for 20 min at room temperature with rotation. Activation and PCR were performed in the same manner as for LOOP-TAG reactions. To calculate *J*-loop, it is assumed that each immobilized DNA target is tagmented at most one time by free Tnp or pA-Tnp, and that both PCR product yield and DNA sequencing library yield are proportional to [Tnp]_free_ (or [pA-Tnp]_free_). This assumption was tested using four [pA-Tnp]_free_ and [Tnp]_free_ concentrations: 2 nM, 0.4 nM, 0.08 nM, 0.016 nM, at 22, 24, and 26 PCR cycles. Based on these results, 2 nM, 0.4 nM, and 0.08 nM Tnp concentrations were chosen at 22 cycles for both annealing-mediated conjugation and antibody-mediated conjugation. It was further confirmed that these conditions yielded proportional nanopore DNA sequencing counts.

#### Conditions ensuring intramolecular tagmentation

An initial estimate was made to assure sufficiently sparse paramagnetic streptavidin bead loading with ∼1,000-bp duplexes such that only intramolecular DNA looping would be detected. M-270 Dynabeads™ streptavidin magnetic beads (Invitrogen #65-601) are reported by the manufacturer to have a mean radius of 1.4 µm so ∼24.63 µm^2^ bead surface (assuming smooth). The stock solution was 6-7×10^8^ beads/mL. Assuming a fully-extended 1110-bp target DNA (∼377 nm) and IgG antibody/pA-Tnp protein complex (∼21 nm), the target DNA/protein conjugate could sweep an area of 0.5 µm^2^, suggesting that target DNA loading should be limited to 49 per bead to prevent interactions between conjugates. For a DNA target with molecular weight 721000 g/mol we then estimated:

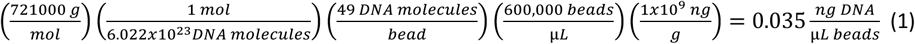

Equation (1) led to the loading of 100 µL magnetic beads with 3 ng target DNA.

Two criteria were then used to confirm that bead loading was sufficiently sparse to assure only intramolecular tagmentation. First, Tnp loaded on one DNA target must tagment only that target and not adjacent DNAs. Second, DNA-loaded beads must not significantly interact with other beads while conjugated Tnp is activated. To validate these assumptions, both DNA-bead combination and DNA target combination experiments were used involving two different DNA targets amplifiable by different PCR primer pairs: one tagmentation-competent DNA target (Target 1) contained a single-stranded annealing terminus (template 1506 amplified by primers 7061/7906) or Dig-labeled terminus (template 1506 amplified by primers 7061/7063) for conjugation with loaded Tnp. The bystander target (Target 3, template 2656 amplified by primers 7036/7034) detectable with distinct diagnostic PCR primers was synthesized with a blunt terminus incapable of loading Tnp. For bead combination experiments, a 25 µL aliquot of beads coupled to Tnp-conjugated Target 1 DNA (template 1506 amplified by primers 7061/7063 or template 1506 amplified by primers 7061/7906) was mixed with a 25 µL aliquot of beads coupled to Tnp-unconjugated Target 3 DNA (template 2656 amplified by primers 7036/7034) to determine whether bead-bead interactions occur allowing Target 3 beads to be tagmented by Target 1 Tnp-conjugated beads. In DNA target combination experiments, 1.5 ng Target 1 DNA Tnp conjugates (template 1506 amplified by primers 7061/7063 or template 1506 amplified by primers 7061/7906 and 1.5 ng bystander Target 3 blunt-ended DNA conjugates (template 2656 amplified by primers 7036/7034) were loaded on the same 100 µL aliquot of M-270 streptavidin magnetic beads to determine if Target 3 bystander DNA strands would be tagmented by Target 1 Tnp conjugates. The LOOP-TAG protocol was then implemented normally. For subsequent PCR analysis, each sample was divided in half. One half of the sample beads were subjected to amplification with tagmentation detection primers for Target 1 DNA and half for Target 3 bystander DNA. The resulting products were subjected to electrophoresis through a 2% agarose gel, stained with ethidium bromide and imaged with a Typhoon imager (Amersham). Target 1 tagmentation in the absence of Target 3 tagmentation demonstrates only intramolecular Target 1 tagmentation, confirming that bead loading and bead interactions are sufficiently sparse.

#### qPCR determination of DNA-bead binding and adapter loading efficiencies

qPCR primers are listed in Supplemental Tables S8-S9. qPCR was used to estimate the adapter-loaded Tnp conjugation efficiency by Tnp terminal annealing. In all cases, qPCR standard curves were created as follows: a 1.33 fmol/µL stock of template 1506 amplified by primers 7061/7063 was diluted to 0.442 fmol/µL and 6 µL of this dilution (2.65 fmol) used as the first point on the standard curve. Alternatively, a 45 µM stock of 7987/7066 adapter was diluted to 0.833 fmol/µL and 6 µL of this dilution used as the first point on the standard curve (5 fmol). Five-fold dilutions in H_2_O were completed for both template 1506 amplified by primers 7061/7063 and adapter 7987/7066 for a total of seven points on each standard curve. Tnp conjugation strategies were followed as described above using 50 µL loaded DNA-beads. Deviations from the above protocol included using 7987/7066 adapter instead of 7065/7066 and resuspension in 100 µL H_2_O instead of Looping Buffer at which point the standard protocol was terminated. Resuspended DNA-beads were incubated for 10 min at 90°C to denature the Tnp and release adapters into solution. After cooling to room temperature, DNA-beads were bound to a magnet. The supernatant was removed for qPCR analysis of 7987/7066 adapter yield. Beads were resuspended in 50 µL H_2_O for quantitating total bead-bound DNA. qPCR (25 µL) was assembled with 12.5 µL SYBR green qPCR mix (QuantaBio PerfeCTa #95071-012), 2 µL forward and reverse primers for full length DNA (10 µM each) or 7987/7066 adapter (5 µM each), 2.5 µL water, and 6 µL DNA-beads or 7987/7066 adapter sample. Cycle temperatures and times were 94°C (3 min), 39 cycles of [95°C (10 s), 57°C (10 s), and 72°C (10 s)]. Conjugation efficiency was determined by calculating the ratio of fmol 7987/7066 adapter to fmol loaded DNA-beads, noting that each Tnp dimer can, in principle, load two adapters: a 2:1 molar adapter:Tnp dimer ratio represents 100% efficiency in assembly. Final estimated tethered and free adapter concentrations were used as constraints in *J*-*loop* calculations. For example, two independent replicates each for two templates gave a mean of 0.0344 fmol adapter (Target 1) and 0.0272 fmol adapter (Target 2) in 50 µL of DNA-beads (0.0172 fmol loaded Tnp and 0.0136 fmol loaded Tnp, respectively) vs. a potential of 0.546 fmol and 0.523 fmol, respectively, bead-bound DNA had 100% assembly been achieved. Therefore, the fraction assembly (*f*) was 0.0315 and 0.0260, respectively, for these preparations of Target 1 and Target 2. The fraction assembly was not determined for each preparation and in subsequent experiments was estimated as a fit parameter.

#### Tagmentation library preparation and nanopore sequencing

Bar-coded DNA libraries were created by PCR amplification of bead-bound tagmentation fragments using anchor primers at the bead terminus of the target DNA and mosaic end A primers at the tagmentation terminus. PCR products from 50 µL reactions were purified using the QIAquick PCR Purification Kit (Qiagen #28104) and resuspended in 25 µL H_2_O. A 3 µL aliquot of purified PCR product was subjected to electrophoresis through a 2% agarose gels to analyze the DNA size distribution after staining with ethidium bromide and imaging on a Typhoon imager (Amersham). Bar-coded tagmentation product libraries were created using ligation sequencing amplicons (Native Barcoding Kit 24 V14 SQK-NBD114.24: Oxford Nanopore #EXP-NBA114) according to a variation of the manufacturer’s protocol (see below) together with the NEBNext^®^ Quick Ligation Module (New England Biolabs #E6056S), NEBNext^®^ Ultra™ II End Repair/dA-Tailing Module (New England Biolabs #E7546S), NEBNext^®^ FFPE DNA Repair Mix (New England Biolabs #M6630S), and blunt/TA Ligase Master Mix (New England Biolabs #M0367L). Resulting bar-coded samples were subjected to nanopore sequencing using an Oxford Nanopore flow cell (#FLO-MIN114) on a GridION (Oxford Nanpore) instrument for 48-72 h. Samples from each bar-coded LOOP-TAG experiment were pooled and run in triplicate on the same flow cell. Variations from the Native Barcoding Kit standard protocol included using an equal volume (12 µL) in the end preparation reaction regardless of DNA concentration (to preserve proportionality of count yields to DNA input), binding DNA products to 2× AMPure XP Beads to preserve short fragments, washing with 80% ethanol rather than Short Fragment Buffer during the Native Barcode Litigation step, and pooling DNA samples after the adapter ligation and clean-up steps using equal volumes of each sample irrespective of concentration.

#### Demonstration of proportionality between input DNA mass and sequencing counts

A DNA library stock of Target 3 DNA was subjected to two-fold dilutions yielding 300 ng, 150 ng, 75 ng, 37.5 ng, 18.75 ng, and 9.375 ng of DNA for input into the Native Barcoding Kit. DNA dilutions were uniquely bar-coded and pooled. Equal volumes of the resulting DNA libraries were subjected to nanopore flow cell sequencing on a GridION instrument and sequencing counts plotted to demonstrate that DNA input mass is proportional to sequencing counts collected over this mass range.

#### Data analysis and modeling

First, min, max, and mean were calculated at each loop length from every triplicate set of counts for each bar-coded LOOP-TAG sample. These min and max values at each loop length were used to plot an envelope of the triplicate measurements. The mean values at each loop length were used to calculate *J-loop* values (calculated separately for each [Tnp]_free_) using equation 2, with the parameter *f* initially taken to be unity:

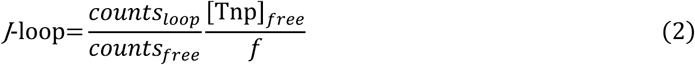

where *counts_loop_* is the intramolecular looping fragment distribution with conjugated Tnp, *counts_free_* is the free Tnp intermolecular fragment distribution, [Tnp]_free_ is the corresponding known concentration of free Tnp, and *f* (fractional assembly) is a parameter from fitting reflecting the efficiency of assembly of adapter-loaded Tnp. This parameter was constrained by experimental values determined from qPCR. All *counts* are lengths from nanopore sequencing of DNA libraries created by PCR amplification of bead-bound tagmentation fragments. Because *J-loop* involves a ratio of terms dependent on Tnp activity, any difference of Tnp activity from 100% cancels in the calculation.

Then min, max, and mean were calculated for *J*-*loop* from the triplicate concentrations. The mean *J*-*loop* values at each loop length were used to construct a cost function for fitting that consisted of a least squares summation between the natural logarithm of the data and the natural logarithm of the model. Fitting was divided into two steps as shown in Supplemental Fig. S1. The first step utilized only the protein-mediated Worm-like Chain (WLC) model as described in (36), based on the work of (37). Equations are reproduced in Supplemental Fig. S1. This first step determined the fractional assembly parameter (*f*) for the efficiency of assembly of adapter-loaded Tnp. In cases where *f* was measured experimentally, the DNA persistence length (*P*) results from the calculation. In other cases, two of the three additional parameters in this step, i.e. *P*, the protein bridge radius (*r*), and the DNA loop kink angle (κ) were fixed. In these cases, *P* was fixed at 50 nm (the conventional low-salt DNA persistence length value) and in all cases κ was fixed at 180° (no kink in the smooth DNA loop). Additionally, this fitting was confined to loop lengths that avoided the anomalous shortest-range hypertagmentation regime and the longest-range regime with higher background signal. In cases where the value of the fractional assembly parameter (*f*) was not experimentally determined, fitted values of the were then fixed for the second step (Supplemental Fig. S1). This second step involved fitting the now corrected (scaled by *f*) mean *J*-loop data over the entirety of loop lengths. A summation model was required to account for the anomalous short-range hypertagmentation regime. For simplicity, this summation model consisted of a generic Lorentzian function (equation in Supplemental Fig. S1), divided into two sides and summed with the protein-mediated WLC model. Specifically, the left-side Lorentzian applies from 0 bp to the bp defining its peak, while the right-side Lorentzian applies from the defined peak of the left-side to longer lengths. The protein-mediated WLC model applies from a minimum length equal to the standard 2*r* (converted from nm to bp using 0.32 nm/bp) to longer lengths. The model parameters of the Lorentzian: peak amplitude (*A*), peak center (*μ*), and half width at half max (γ) were heavily constrained by the data, resulting in excellent fits. For the left-side Lorentzian, *A* was fixed at the *J*-loop maximum, while *μ* and γ were constrained (Supplemental Table S10). For the right-side Lorentzian, *A* was fixed at the *J*-*loop* maximum obtained with 10 nM Nhp6A (if it was higher, otherwise the normal maximum *J*-loop), *μ* was fixed at 0 bp, and γ was constrained (Supplemental Table S10). Likewise, in experiments where *f* was not explicitly measured, the protein-mediated WLC model was constrained: *P* was fixed at 50 nm, *r* was constrained, and κ was fixed at 180°. The cost function for minimizing the four variables to be fitted included two additional penalties. One penalty ensured that the two sides of the Lorentzian met at the same value and the other penalty created agreement between the summation model minimum and the protein bridge radius. Final fitted values of all parameters are given in Supplemental Table S10. Area under the curve calculations were then made from either mean experimental data values or model components using the trapezoidal approximation.

## Results

### A massively-parallel protein-mediated DNA looping assay

DNA LOOP-TAG is shown conceptually in Fig. 1C-D. Terminally-biotinylated ∼1,000-bp duplex DNA sequences are immobilized at low density on streptavidin-modified magnetic beads. Bead tethering at a sparse loading density ensures that only intramolecular looping events are detected as tethered DNAs cannot interact with one another. The distal terminus of each DNA is conjugated with Tn5 transposase (Tnp). Upon activation with Mg^2+^ ions, intramolecular DNA looping allows tagmentation with single-hit kinetics. Shorter bead-bound tagged DNA fragments correspond to larger DNA loops. The distribution of lengths of resulting bead-bound tagged fragments reports the probability of each Tnp-mediated DNA loop size. PCR amplification of these bead-bound fragments, bar-coded library synthesis, and nanopore-based sequencing, allow highly-reproducible quantitation of loop length probabilities through millions of fragment reads (*counts_loop_*). We have found nanopore sequencing-based determination of DNA length distributions to be the most reliable and accurate approach available. To convert this probability distribution to a *J*-looping distribution, an experiment is performed with an identical bead-tethered ∼1,000-bp DNA preparation lacking conjugated Tnp but exposed to a known concentration of free Tnp under the same experimental conditions for the same time (*counts_free_*). The ratio of the intramolecular looping fragment distribution with conjugated Tnp (*counts_loop_*) to the free Tnp fragment distribution (*counts_free_*), multiplied by [Tnp]_free_ gives a *J*-*loop* value for every Tnp-mediated fragment length of the ∼1,000-bp tethered DNA at single bp resolution.

We first developed a Tnp conjugation strategy to minimize the size and flexibility of the tethered Tnp (Fig. 2A). Here the distal terminus of the ∼1,000-bp tethered target DNA carries a short-single-stranded sequence that permits annealing to one mosaic A adapter in a Tnp complex loaded with two such adapters. This Tnp protein complex is deliberately compact with a diameter less than 13 nm and a short linker. This configuration is intended to make intramolecular tagmentation largely dependent on DNA physical properties, though flexibility is present in the tri(ethylene glycol) linker appending the distal DNA single strand to the tethered ∼1,000-bp DNA. Conversion of intramolecular tagmentation events to *J-loop* values depends on a control experiment (Fig. 2B) with a known concentration of free Tnp exposed to the same tethered construct under the same conditions for the same time.

**Fig. 2.**
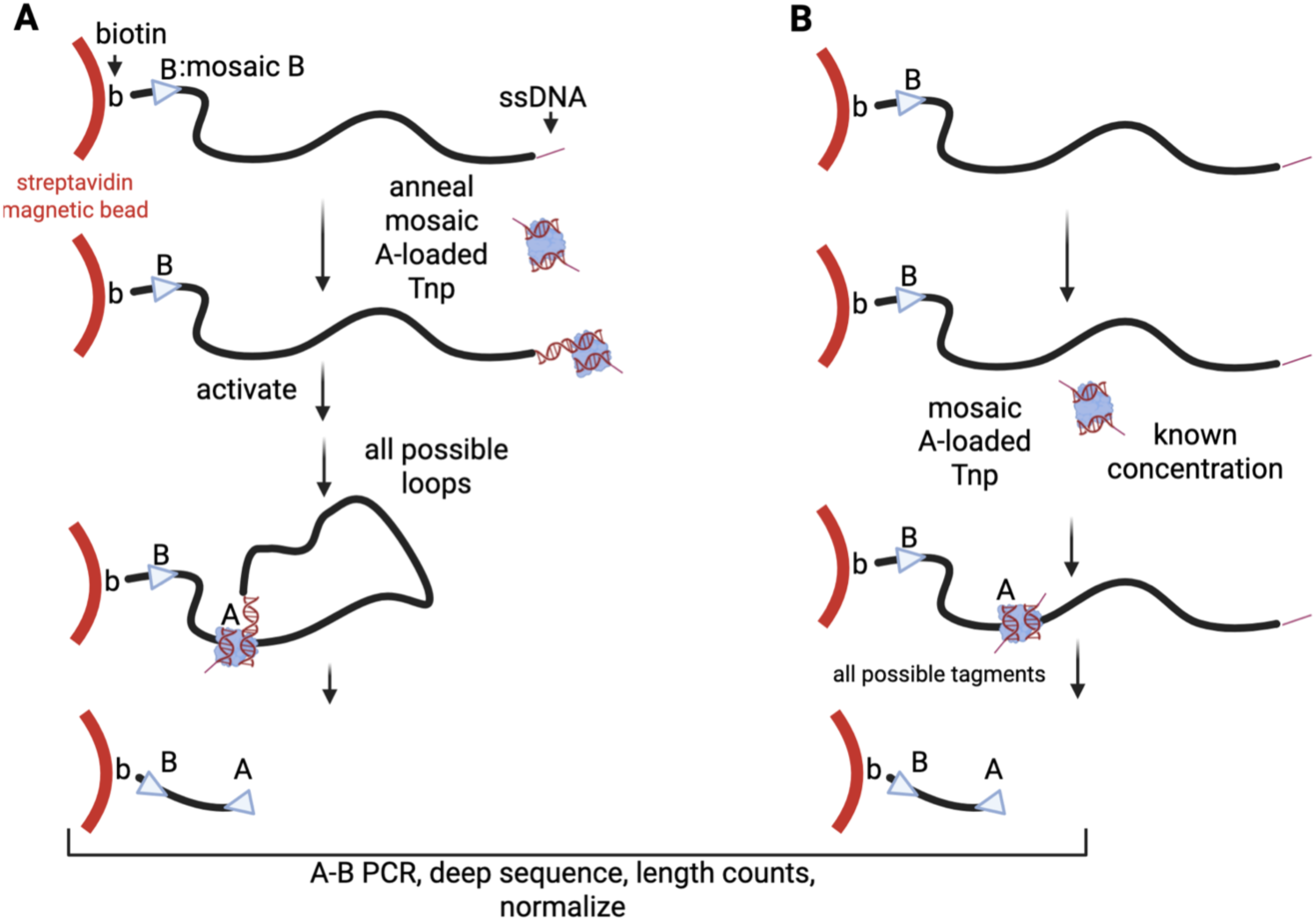
Details of intramolecular protein-mediated DNA looping assay based on a small and inflexible looping protein. Conjugation involved assembly of a complex of Tn5 Tnp loaded with mosaic A DNA adapters, one of which is annealed to a single-stranded complement at the distal terminus of the ∼1,000-bp bead-immobilized DNA. Tagmentation by (A) tethered Tnp, or (B) a known concentration of free Tnp. Looping constructs (∼1,000 bp) are biotinylated (b) on one terminus and conjugated at low density to streptavidin-coated magnetic beads. Looping constructs contain a mosaic B sequence adjacent to the bead. Tagmentation by intramolecular DNA looping is initiated by addition of Mg2+ ions. In (B), free Tnp loaded with mosaic A adapters is allowed to tagment in the presence of Mg2+ ions for the same time as in (A). The resulting bead-bound tagments are amplified by PCR primers complementary to mosaic A and B sequences, and the resulting DNA fragment library is subjected to nanopore-based deep sequencing to determine counts for each tagment length. After correcting for conjugate assembly efficiency, counts obtained from libraries produced by intramolecular DNA looping in (A) are divided by corresponding counts obtained in (B). The quotient for each length multiplied by the free transposase concentration used in (B) defines *J*-loop for each loop length.

We then developed a second Tnp conjugation strategy (Fig. 3A), where the ∼1,000-bp target DNA carries a terminal digoxygenin (dig) modification, allowing binding by an anti-dig IgG antibody which, in turn, can be bound by a commercially-available Protein A-Tn5 Tnp fusion protein (pA-Tnp) loaded with mosaic A adapter DNA oligonucleotide duplexes. This protein assembly is deliberately larger (estimated diameter ∼21 nm) and flexible, intended to increase the looping probability by reducing required DNA bending. In both cases, tagmentation occurs within the ∼1,000-bp bead-bound DNA upon intramolecular looping. As in the compact design, conversion to J-loop values requires an experiment with free Tnp (Fig. 3B).

**Fig. 3.**
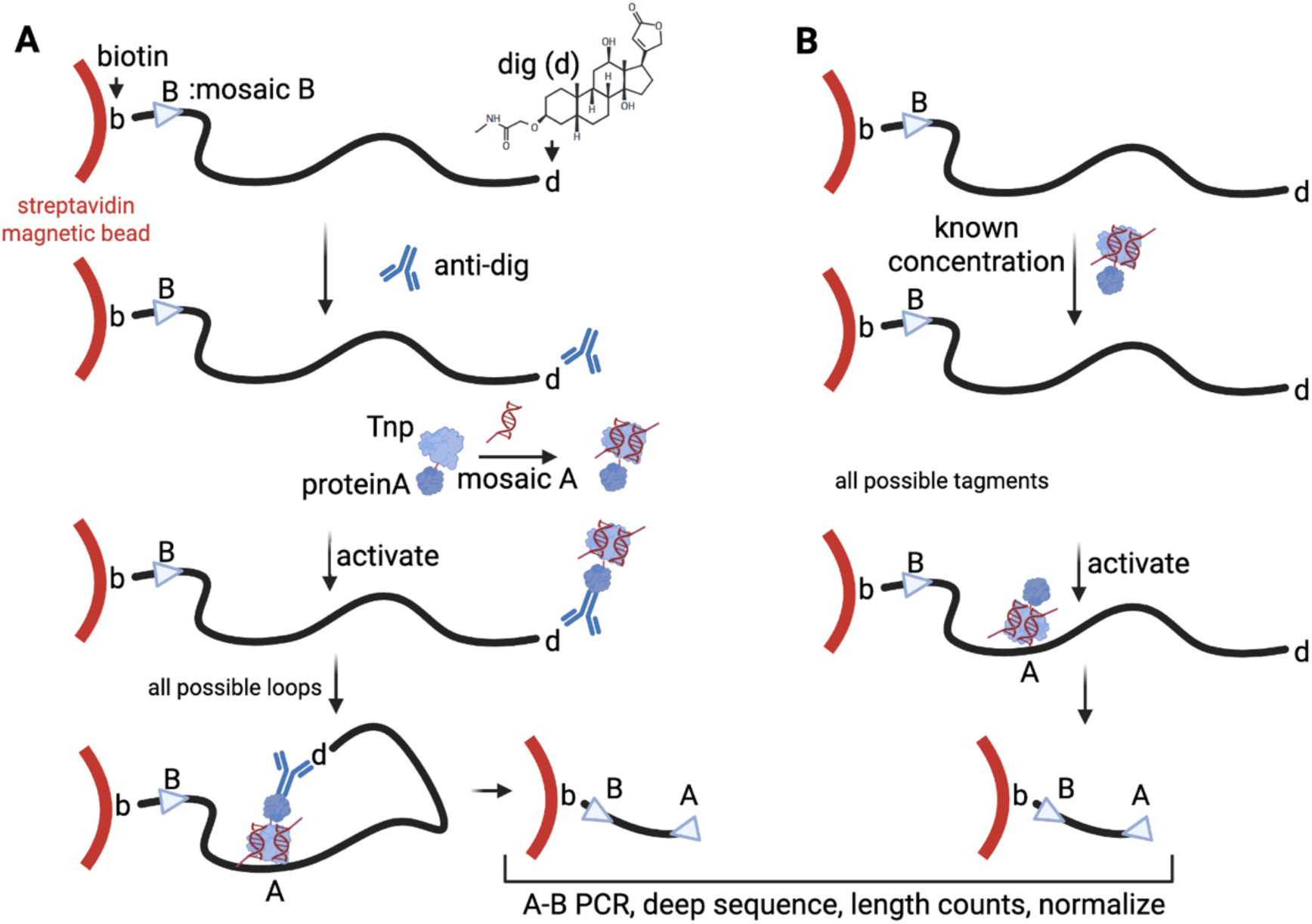
Details of intramolecular protein-mediated DNA looping assay based on a large and flexible looping protein complex. The distal terminus of the looping construct caries a digoxygenin (dig, d) epitope, recognized by an anti-dig IgG monoclonal antibody. Subsequent binding of a Tn5 Tnp-protein A fusion protein (pA-Tnp) loaded with two copies of a mosaic A adapter sequence creates the mature looping construct with flexibly-tethered Tnp. Tagmentation by (A) flexibly-tethered Tnp, or (B) a known concentration of free pA-Tnp. Details are as in Fig. 2.

### Validation of experimental systems

Quantitative success of the LOOP-TAG approach depends on multiple assumptions that were validated. Adapter loading into Tnp proteins must be efficient. This assumption is validated in Supplemental Fig. S2 where efficient DNA duplex adapter loading into pA-Tnp and Tnp proteins is shown as detected in a native polyacrylamide gel (Supplemental Fig. S2A), a native blue polyacrylamide gradient gel (Supplemental Fig. S2B), vs. a denaturing polyacrylamide gel (Supplemental Fig. S2C). Tnp activation must be controllable by Mg^2+^ addition or chelation and Tnp conjugation must be predominantly at the distal terminus of the tethered ∼1,000-bp DNA, with only low levels of Tnp adhering to the body of the ∼1,000-bp DNA after assembly. These assumptions are validated for annealed Tnp conjugates in Fig.4 and for antibody-mediated Tnp conjugates in Fig. 5. In both cases, it is shown that tagmentation is only observed in the presence of Mg^2+^ ions and when the bead-tethered DNA carries an annealed terminal Tnp. Tethering of ∼1,000-bp DNA molecules must be sufficiently sparse on streptavidin magnetic beads that looping results only in intramolecular tagmentation. This assumption is validated for annealed Tnp conjugates in Fig. 4B and for antibody-mediated Tnp conjugates in Fig. 5B. In both cases, intramolecular tagmentation is demonstrated to occur without intermolecular tagmentation by including an un-conjugated tethered DNA strand with a different sequence, enabling differential detection by PCR. Sequencing libraries and resulting nanopore sequencing counts must faithfully reflect original numbers of each length of bead-bound DNA fragment after tagmentation. This assumption is validated for annealed Tnp conjugates and antibody-mediated Tnp conjugates by demonstrating proportionality of tagmentation fragment counts to [Tnp]_free_ (Supplemental Fig. S3A-D), and proportionality of DNA input to total sequencing counts (Supplemental Fig. S3E). Finally, it is necessary to estimate the efficiency of assembly of bead-bound, mosaic adapter-loaded ∼1,000-bp DNA-Tnp conjugates. Such estimates were achieved by qPCR determination of bead-associated mosaic A adapters after conjugate assembly on bead-bound DNA, indicating typical overall assembly efficiencies of ∼3%, described in Material and Methods. The calculated fractional assembly parameter (*f*) for one preparation of Target 1 was explicitly determined to be 0.0315, leading to a fitted persistence length (*P*) of 52.8 nm (Fig. 6E). Because this value is near the canonical 50 nm persistence length for duplex DNA, for simplicity, subsequent analyses assumed a persistence length of 50 nm without performing qPCR-based conjugate assembly efficacy determination in each case. Instead, the fractional assembly parameter (which scales the *J*-loop data) was fitted to achieve a 50 nm persistence length. It should be emphasized that the LOOP-TAG methodology combined with qPCR determination of conjugate assembly efficiency allows explicit determination of the DNA persistence length of the test sequence.

**Fig. 4.**
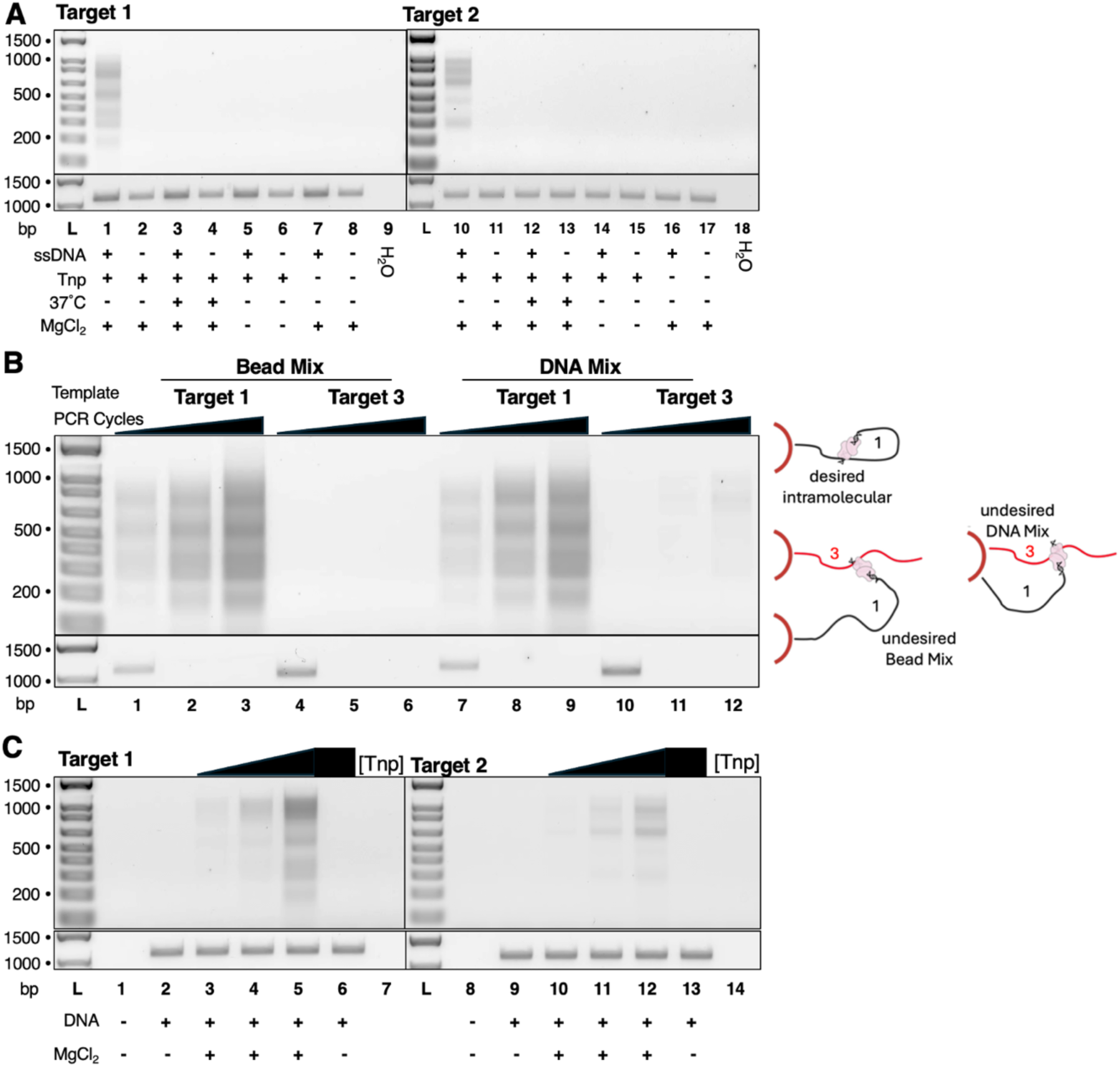
Validation of key LOOP-TAG parameters for Tnp-annealed conjugates by 0.8% agarose gel analysis of PCR products. DNA ladder (L) sizes (bp) are shown (left). See Methods for full experimental details. A. Mosaic A adapter (7065/7901)-loaded Tnp (lanes 1-6 and 10-15) was incubated with the indicated magnetic bead-immobilized DNA targets with (lanes 1, 3, 5, 7, 10, 12, 14, 16) or without (lanes 2, 4, 6, 8, 11, 13, 15, 17) ssDNA overhangs for conjugation by annealing. The magnetic bead complex was washed to remove non-specifically bound Tnp. Where indicated, beads were incubated at 37°C to remove annealed Tnp complex (lanes 3-4 and 12-13). Tagmentation was initiated with MgCl2 (lanes 1-4, 7-8, 11-13, and 16-17). Template-free PCR controls are shown (H2O). B. Validation of intramolecular tagmentation (schematic at right). Bead mix samples (lanes 1-6) combined equivalent amounts of Target 1 (ssDNA overhang) and Target 3 (blunt) DNA-bound magnetic beads prior to Tnp exposure. DNA mix samples (lanes 7-12) loaded equal amounts of Target 1 (ssDNA overhang) and Target 3 (blunt) DNA onto the same magnetic beads. Tagmented products from bead mix and DNA mix were interrogated by PCR for intramolecular (Target 1) and intermolecular (Target 3) tagmentation at 22-26 PCR cycles (top panel). Intact Target 1 (lanes 1 and 7) and Target 3 (lanes 4 and 10) markers were amplified for 15 cycles (bottom panel) C. Tagmentation of bead-bound Target 1 (lanes 2-6) and Target 2 (lanes 9-12) DNA by 0.08 nM, 0.4 nM, and 2 nM free Tnp. Bead-only (lanes 1 and 8) and –MgCl2 (lanes 6 and 13) served as controls.

**Fig. 5.**
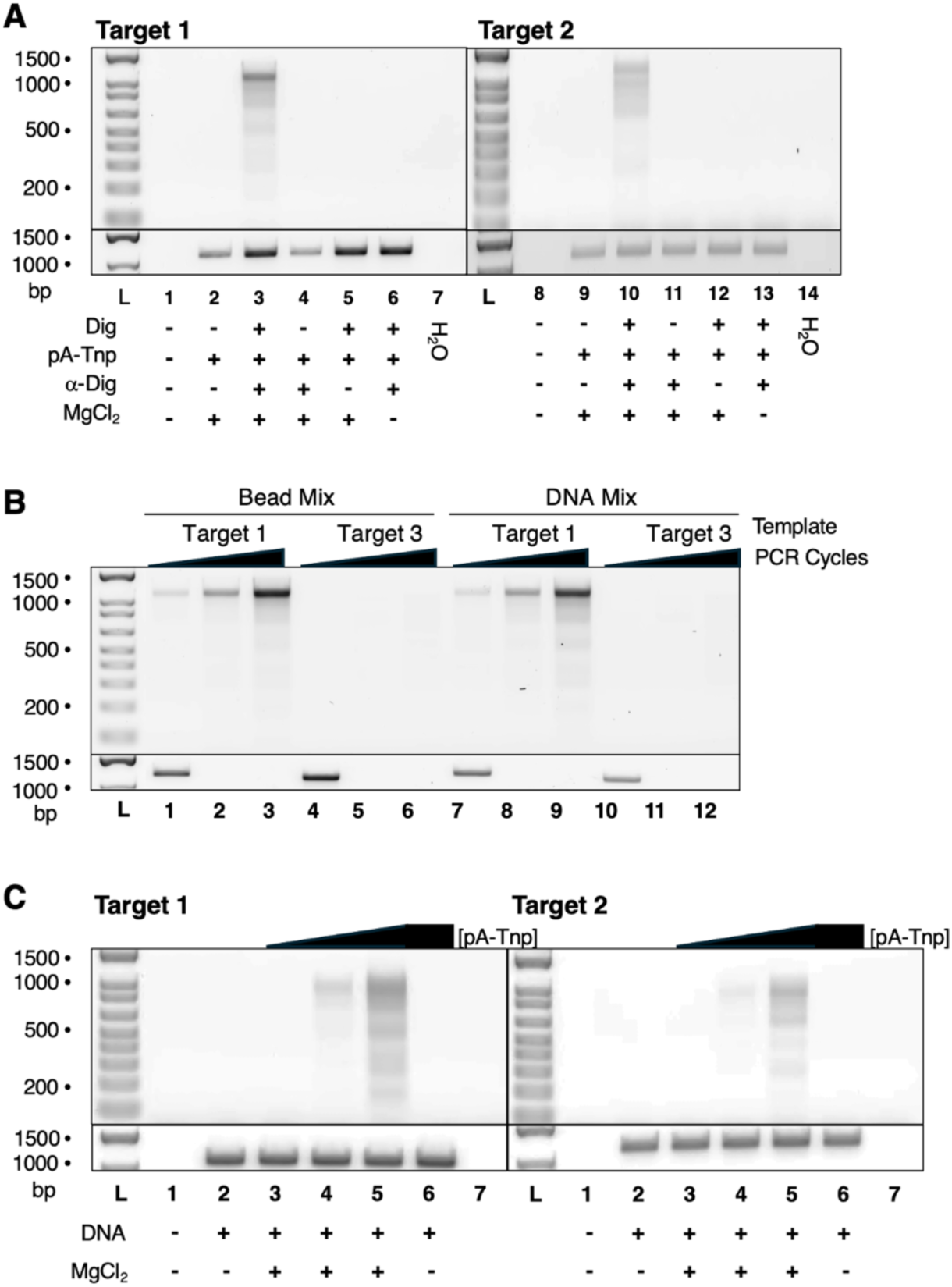
Validation of key LOOP-TAG parameters for antibody-mediated Tnp conjugates by 2% agarose gel analysis of PCR products. DNA marker (L) sizes are shown (left). See Supplemental Methods for full experimental details. A. The indicated magnetic bead-immobilized DNA targets with a terminal digoxigenin (Dig) tag (lanes 3, 5-6,10,12-13), blunt terminus (lanes 2, 4, 9, and 11) or beads alone (lanes 1, 8) were incubated with an α-Dig antibody followed by addition of mosaic A adapter (7065/7066)-loaded pA-Tnp (lanes 1-6, 10-15). Beads were washed to remove non-specifically-associated pA-Tnp. Tagmentation was initiated with MgCl2 (lanes 2-5, 9-13) followed by pA-Tnp inactivation. Template-free PCR controls are shown (H2O). B. Validation of intramolecular tagmentation (schematic at right). Bead mix samples (lanes 1-6) combined equivalent amounts of Target 1 (Dig) and Target 3 (blunt) magnetic beads prior to exposure to α-Dig antibody and pA-Tnp. DNA mix samples (lanes 7-12) loaded equal amounts of Target 1 (Dig) and Target 3 (blunt) DNA onto the same magnetic beads. Tagmented products from bead mix and DNA mix samples were interrogated by PCR for intramolecular (Target 1) vs. intermolecular (Target 3) tagmentation at PCR cycles 22-26 (top panel). Intact Target 1 (lanes 1,7) and Target 3 (lanes 4,10) markers were amplified for 15 cycles (bottom panel). Interpretation diagram shown to right. C. Tagmentation of bead-bound Target 1 (lanes 2-6, left) and Target 2 (lanes 2-6, right) DNA by 0.08 nM, 0.4 nM, and 2 nM free Tnp. Bead-only (lanes 1 and 8) and –MgCl2 (lanes 6 and 13) are controls.

**Fig. 6.**
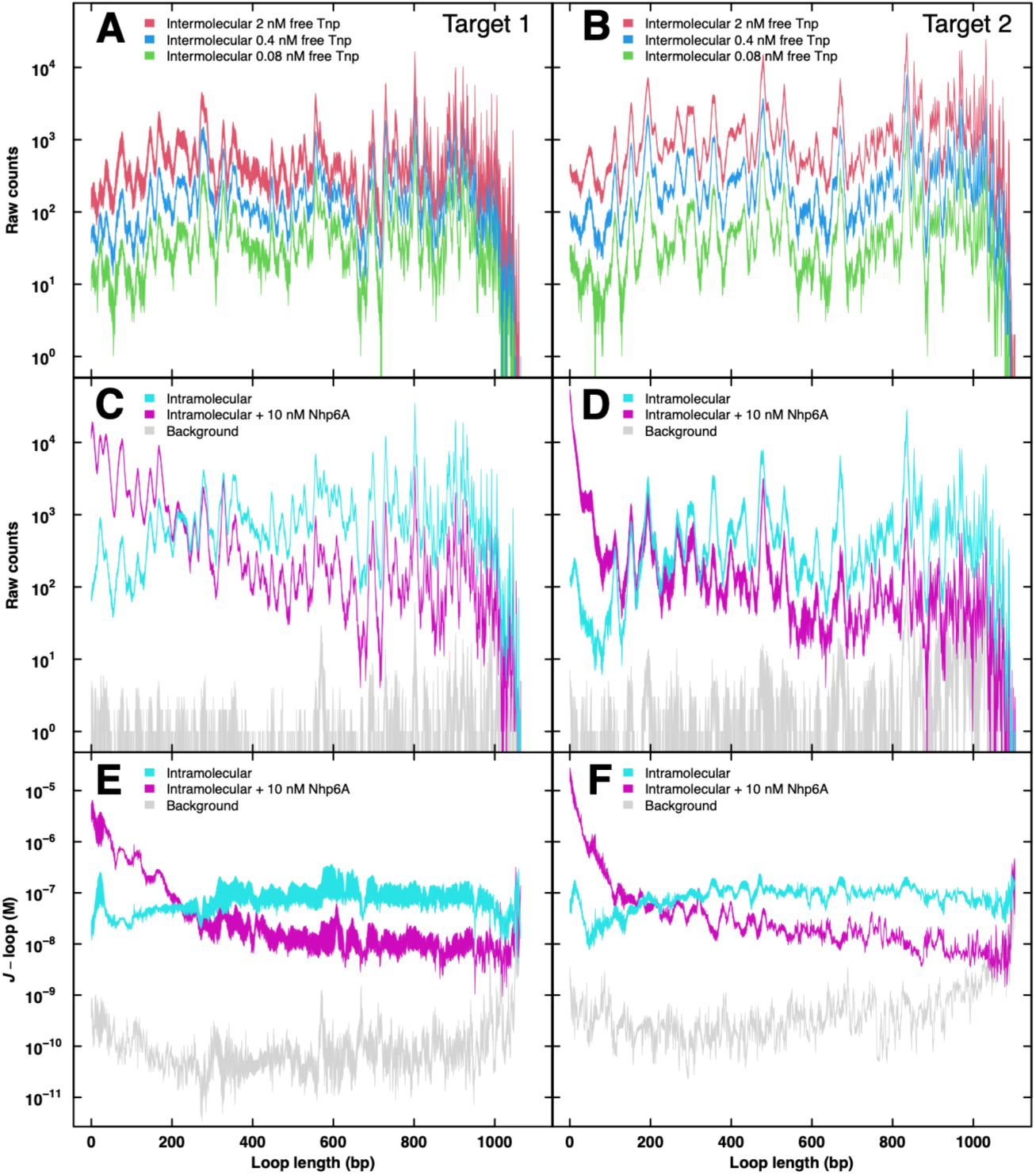
LOOP-TAG. Loop probabilities for the indicated target DNAs determined by LOOP-TAG with an annealed terminal Tnp. A-B. Length-dependent bead-bound target DNA tagmentation PCR-amplified fragment counts from nanopore sequencing after exposure to the indicated [Tnp]free. C-D. Length-dependent bead-bound target DNA tagmentation PCR-amplified fragment counts from nanopore sequencing after intramolecular looping of Tnp-annealed conjugates in the absence (cyan) or presence (magenta) of 10 nM Nhp6A protein. E-F. The length-dependent *J*-looping values in the absence (cyan) or presence (magenta) of 10 nM Nhp6A protein were either directly determined from Equation 2 using a measured value of *f* (E) or appropriately scaled by a fitted value of *f* to yield *P* = 50 nm (F). Background signal (counts after thermal removal of annealed Tnp) are shown in grey (C-F). In all cases, traces reflect the envelope of experimental triplicate measurements.

A known feature of Tn5 Tnp tagmentation is strong DNA sequence preference (38). Our approach to determining *J*-looping accommodates this known bias. As described in Materials and Methods equation 1, *J*-loop estimates adjusted for assembly efficiency are calculated as ratios of counts for all fragment lengths generated by *intra*molecular tagmentation vs. counts for all fragment lengths generated by *inter*molecular tagmentation in the presence of free Tnp at known concentrations. Thus, any DNA sequence-dependent bias in tagmentation or PCR amplification efficiency is present both in the numerator and denominator of the *J*-loop ratio for each fragment length, normalizing the result.

### *J*-looping deduced for loops involving small and rigid vs. large and flexible Tnp conjugates

Having validated necessary assumptions for LOOP-TAG we undertook LOOP-TAG analysis of ∼1,000-bp DNA templates 1 and 2 using both annealing and antibody-mediated terminal Tnp conjugation strategies. The striking results for the compact annealed Tnp case are shown in Fig. 6, and for the large antibody-mediated case in Fig. 7. Length frequency data from nanopore sequencing of PCR products of calibration cleavage products of DNA Targets 1 and 2 exposed to three [Tnp]_free_ are shown in Fig. 6AB (compact Tnp conjugate) and Fig. 7AB (large antibody-mediated Tnp conjugate).

**Fig. 7.**
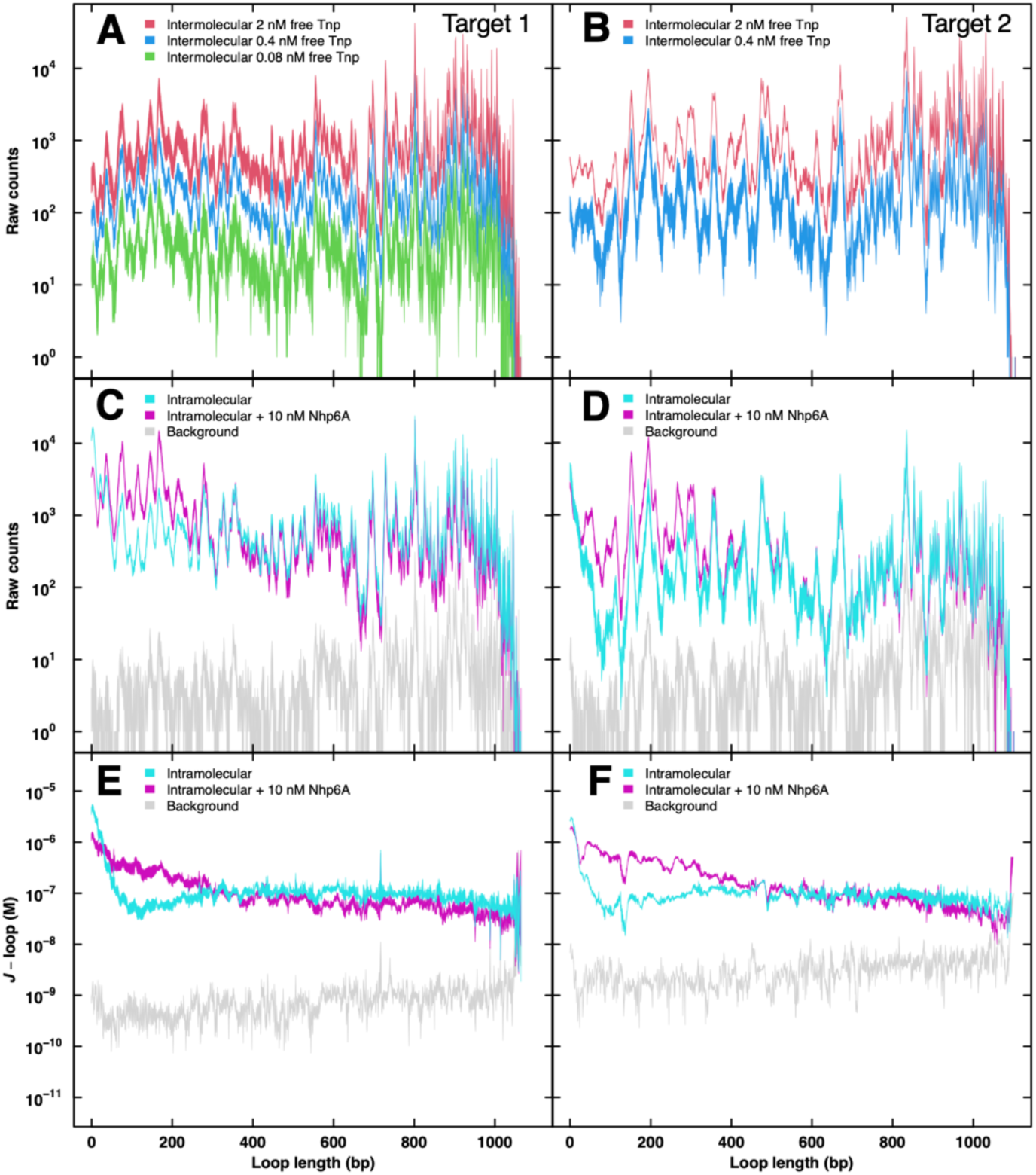
LOOP-TAG. Loop probabilities for the indicated target DNAs determined by LOOP-TAG with an antibody-linked Tnp. A-B: Length-dependent bead-bound target DNA tagmentation showing PCR-amplified fragment counts from nanopore sequencing after exposure to the indicated [Tnp]free. C-D. Length-dependent bead-bound target DNA tagmentation PCR-amplified fragment counts from nanopore sequencing after intramolecular looping of antibody-linked Tnp conjugates in the absence (cyan) or presence (magenta) of 10 nM Nhp6A protein. E-F. Calculated length-dependent *J*-looping in the absence (cyan) or presence (magenta) of 10 nM Nhp6A protein. Background signal (counts after thermal removal of annealed Tnp) are shown in grey (C-F). In all cases, traces reflect the envelope of experimental triplicate measurements.

Length frequency data from nanopore sequencing of PCR products of bead-bound tagmentation products from intramolecular self-cleavage of DNA Targets 1 and 2 for compact annealed Tnp conjugates are shown in Fig. 6CD and for antibody-mediated conjugates in Fig. 7CD. The resulting *J*-loop estimates are then plotted for annealed and antibody-mediated Tnp conjugates in Fig. 6F and Fig. 7EF, respectively, with Fig. 6E directly taking into account experimental efficiency of assembly of adapter-loaded conjugated Tnp. Based on a conventional assumption of the DNA persistence length of 50 nm in low-salt buffer, an initial fit using the protein-mediated Worm-like Chain Model (WLC) (36) was used to estimate the parameter for adapter-loading efficiency (*f*) in subsequent cases. Uniquely, LOOP-TAG generates *J-*loop at single bp resolution for all DNA fragments terminating in Tnp. From Fig. 6 involving the small and compact annealed Tnp assembly, several key results are evident. The first is the expected high sequence-specificity of tagmentation.

The second and most obvious feature is a striking enrichment of very short loops consistent with an artifact that we attribute to a tagmentation regime in which tethered Tnp folds back via local DNA deformation and the flexible tri(ethylene glycol) linker separating the Tnp from the ∼1,000-bp target (Fig. 6EF, cyan). An even more pronounced anomaly of this kind is seen for antibody-mediated Tnp conjugates (Fig. 7EF, cyan). We modeled this anomalous short-range intramolecular cleavage propensity by fitting a generic Lorentzian function as described in Supplemental Fig. S1 such that the *J*-loop data reflect the sum of the anomalous local tagmentation function and the underlying WLC behavior of protein-mediated intramolecular DNA looping. Results of this fitting procedure are shown in Fig. 8 for both annealed and antibody-based Tnp conjugates. These results give a most likely looping probability of ∼290 bp, notable in comparison to the value of ∼450 bp for T4 DNA ligase cyclization (1). We interpret this as evidence that the Tnp protein conjugates participate as part of the loop, reducing required DNA deformation.

**Fig. 8.**
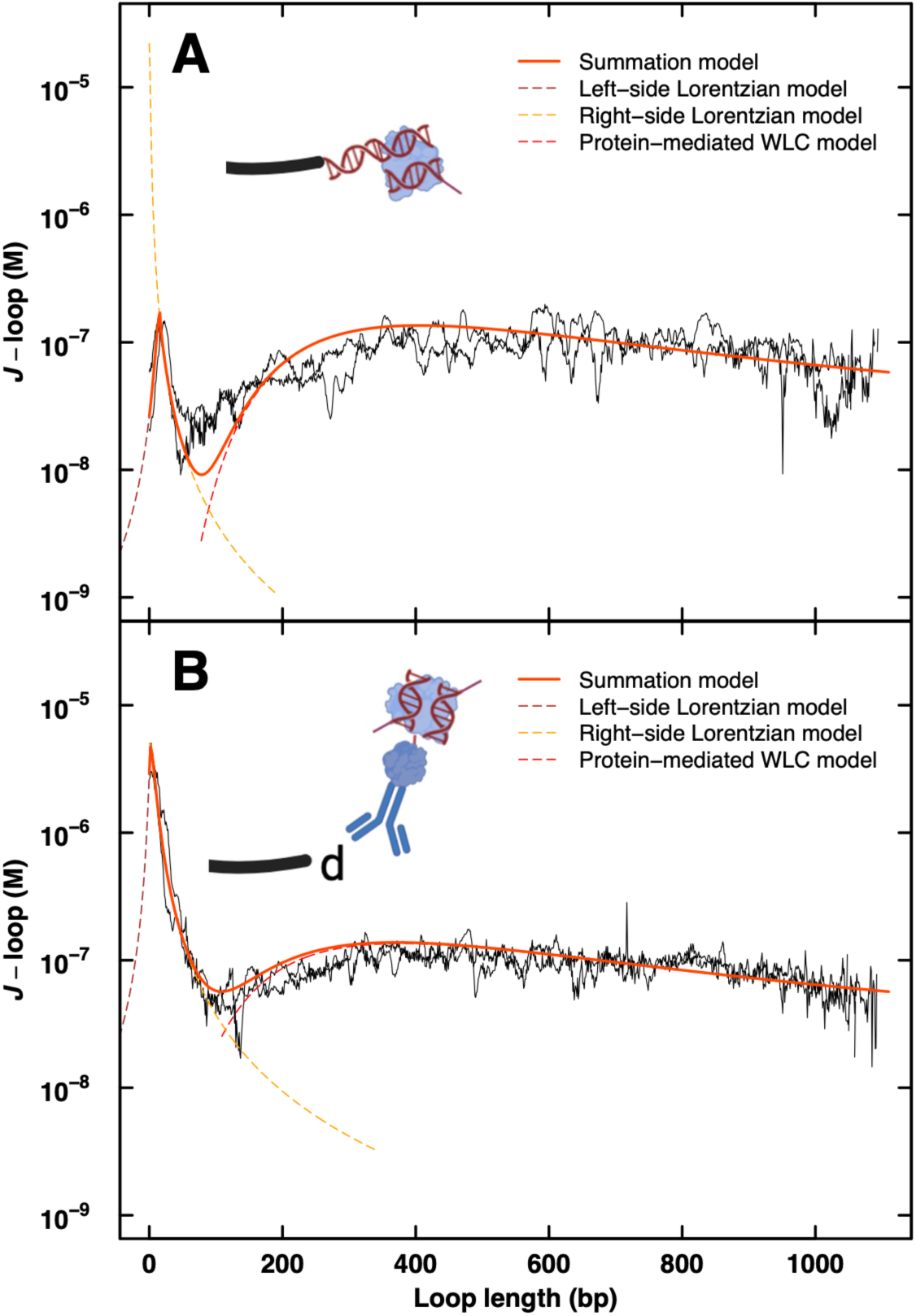
Effect of Tnp complex size and conjugation configuration on anomalous local tagmentation. Full details of data analysis and modeling are described in Methods. Corrected (scaled by *f*) mean *J*-loop data for DNA Targets 1 and 2 are shown in black. A summation model was used for fitting these data to account for the anomalous short-range tagmentation regime. This summation model (bold orange-red) implemented a generic Lorentzian function divided into two sides (left-side, brown; right-side, orange) summed with the protein-mediated WLC model (red). Model fit parameters appear in Supplemental Table S10.

We further reasoned that analysis of the anomalous *J*-loop contributions from local tagmentation can be used to derive estimates for the effective dimensions of the Tnp complexes bridging the DNA loop. The loop lengths corresponding to minimum values of the summation model (78 bp and 109 bp for annealed and antibody-based Tnp conjugates, respectively) suggest that the local tagmentation anomalies occur only within distances of ∼25 nm and ∼35 nm, respectively. The larger antibody-based Tnp conjugate has greater reach as expected. However, the position of the anomalous peak (16 bp and 3 bp for annealed and antibody-based Tnp conjugates, respectively) suggests that the small and compact annealed Tnp assembly suffers from a restrictive tether that limits fold-back geometries relative to the larger antibody-based Tnp conjugate. Anomalous local hypertagmentation can be quantified by calculating the excess area under the summation model curve relative to the protein-mediated WLC model curve. Consistent with intuition, this excess area is 17-fold larger for the antibody-based Tnp conjugate than for the compact annealed Tnp conjugate.

Leaving aside the anomalous hypertagmentation adjacent to the conjugated Tnp protein, the shapes of the fit curves in Fig. 6EF (cyan) and Fig. 7EF (cyan) show a ∼10-fold range in *J-loop*, values, just as anticipated for this portion of the *J-loop* curve according to the wormlike chain polymer model.

### Quantitation of DNA apparent flexibility enhancement by an architectural DNA binding protein

Because the LOOP-TAG assay provides a quantitative approach for massively parallel determination of *J*-loop values for thousands of DNA loop lengths, we applied the method to a system where apparent DNA flexibility can be manipulated. An ideal example is provided by DNA looping in the presence of the high mobility group B (HMGB) yeast family member Nhp6A (39,40). This architectural DNA binding protein sharply kinks DNA (∼90°) by engaging the minor groove with little sequence-specificity. Random binding and unbinding of DNA by Nhp6A have the effect of creating an ensemble of more compact DNA shapes, theoretically enhancing the probability of small DNA loops detectable by the tagmentation fragment size distributions in LOOP-TAG.

We first tested free Tnp activity on bead-tethered DNA in the presence of different concentrations of Nhp6A (Supplemental Fig. S4). Minimal effects were observed so for simplicity subsequent *J*-loop calculations were performed using normalization to free Tnp without Nhp6A. The results of such LOOP-TAG experiments involving 10 nM Nhp6A are shown for the annealed Tnp conjugate in Fig. 6EF and for the antibody-mediated conjugate in Fig. 7EF. In both cases the presence of 10 nM Nhp6A strikingly alters the LOOP-TAG fragment distribution to favor small DNA loops, consistent with random DNA kinking. This redistribution occurs at the expense of larger DNA loops, an effect that is most obvious in Fig. 6 for both DNA Targets 1 and 2. The local tagmentation anomaly becomes even more pronounced. Fig. 6 EF indicate the most probable anomalous loop length in the presence of Nhp6A is now 3 bp. This is followed by a steady decrease in *J*-loop values until a plateau in the 200–400 bp region, where a peak had been observed for naked DNA, followed by further decline at longer loop lengths. We considered this result from the perspective of the number of successful tagmentation events that occur in a given period of time. If this number is constant in the presence of 10 nM Nhp6A, then the increased rate of successful short-loop tagmentations will be accompanied by a corresponding decreased rate of successful long-loop tagmentations. However, if Nhp6A enhances overall tagmentation rate, increased successful short-loop tagmentations may not result in a proportional reduction in long-loop tagmentations. This latter condition appears to apply to the antibody-mediated Tnp conjugate as shown in Fig. 7EF suggesting the possibility of cooperative Tnp-Nhp6A interactions for this conjugate. These data exemplify the ability of the LOOP-TAG approach to detect and quantitate altered DNA flexibility.

## Discussion

Classical measurement of DNA flexibility or static curvature *in vitro* involves estimation of *J*-factors (effective local molar concentration of DNA fragment termini) based on T4 DNA ligase cyclization kinetics under a set of restrictive assumptions and conditions (15). More recent assays to characterize relative DNA flexibility or static curvature involve FRET detection of cyclization for surface-immobilized DNA fragments with long cohesive single-stranded termini (16,21,22). The shared limitation of these methods is that only one DNA fragment length can be evaluated at a time. T4 DNA ligase-mediated cyclization kinetics experiments are particularly tedious, requiring DNA radiolabeling and time-course measurements involving quantitation of polyacrylamide gel autoradiograms. We hypothesized that quantitative massively-parallel DNA flexibility measurements could be achieved *in vitro* by analyzing the distribution of protein-mediated loop lengths for many copies of a single long (e.g. ∼1,000-bp) bead-immobilized DNA sequence conjugated with a terminal tagmentation enzyme. We selected commercially-available promiscuous Tn5 Tnp variants that cleave target DNA and simultaneously insert terminal adapter sequences. This allows rapid PCR amplification of the population of resulting bead-immobilized tagged DNA fragments and conversion to a library for inexpensive nanopore sequencing at a depth of many millions of reads, thus allowing high-resolution quantitation of the DNA loop length probability distribution for all loops adopted by a ∼1,000-bp test sequence. When combined with data obtained by the tagmentation of the same immobilized DNA sequence with a known concentration of free Tnp, a simple calculation yields *J*-loop estimates for each DNA length.

The LOOP-TAG method depends on a number of key assumptions that were addressed in this work. The most important considerations involve showing that Tnp conjugates tagment only intramolecularly, that original DNA fragment length distributions are faithfully retained after PCR, nanopore library preparation, and nanopore sequencing, and that the efficiency of tethered conjugate assembly can be estimated accurately. We confirmed that these and other important assumptions are valid.

We then applied LOOP-TAG to analyze DNA flexibility in the context of two different Tnp conjugation formats, with two different ∼1,000-bp DNA sequences, and in the absence or presence of the yeast sequence non-specific DNA kinking protein, Nhp6A. In each case, quantitative measurement of DNA flexibility is achieved in terms of most probable loop size and *J*-loop estimates for all DNA lengths. Reproducibility is shown to be extremely high. It would be desirable to quantitate the effects of Nhp6A on the persistence length, but this has proven challenging even for T4 DNA ligase cyclization experiments (41). In LOOP-TAG, the accentuated local hypertagmentation anomaly prevents analysis beyond the obvious qualitative effects that favor smaller protein-mediated DNA loops at the expense of larger loops, as expected for a DNA-kinking architectural DNA-binding protein and single-hit Tnp kinetics.

While achieving many of its goals, the present implementation of LOOP-TAG suffers from limitations. The method and its *J*-loop determinations are not directly comparable to T4 DNA ligase cyclization kinetics because LOOP-TAG involves protein-mediated DNA loops. We argue that such loops are more biologically relevant, but the measured *J*-loop values are dependent on the size and flexibility of the conjugated proteins that participate in the loop, enhancing loop probability by reducing the required degree of DNA deformation (1). In the current setting, both antibody-protein A-mediated conjugation and direct Tnp annealing onto ∼1,000-bp DNA termini lead to Tnp complexes with various degrees of local flexibility. This flexibility generates terminus-proximal hypertagmentation that creates an excess short-fragment anomaly not due to DNA looping. This, together with single bonds allowing conjugate rotation, obscures face-of-the-helix effects that are prominent as length-dependent oscillations in J-factor values from ligation kinetics experiments of fully duplex DNA with short single-stranded cohesive termini. Fourier analysis of our data did not reveal reproducible evidence for oscillation of *J*-loop values with DNA helical repeat. The absence of obvious helical repeat-dependence in looping may also be attributed to the profound sequence dependence of Tn5 Tnp, clearly evident from our data. We had anticipated that because transformation of sequencing counts to *J*-loop values involves division of intramolecular looping tagmentation counts by free Tnp tagmentation counts from the same immobilized constructs, effects of Tnp sequence-specificity and sequence effects on PCR efficiency would cancel, leaving smooth *J*-loop curves. While we show that such smoothing reproducibly occurs to some extent, Tnp sequence specificity remains a feature of computed *J*-loop curves.

We had also initially imagined that LOOP-TAG data might shed light on the controversial question of DNA flexibility at lengths below ∼150 bp (one persistence length). As we have reviewed (1), DNA flexibility in this regime has been the subject of debate. Unfortunately, the protein-mediated nature of the LOOP-TAG protocol introduces Tnp-proximal hypertagmentation artifacts that obscure DNA flexibility for the shortest loops of interest. Most evident for the case of Tnp conjugation via large and flexible antibody-protein A bridging, the Tnp conjugate presumably can fold back and tagment locally in a DNA loop-independent manner that results in spuriously high *J*-looping values for the shortest loop lengths. We show that even conjugates based on direct Tnp annealing onto a terminal single-stranded DNA complement at the distal terminus of the ∼1,000-bp DNA target show an excess local tagmentation probability artifact, an effect we attribute to Tnp wrapping and bending of local DNA and the short, flexible tri(ethylene glycol) linker. Interestingly, the presence of architectural protein Nhp6A modifies this local tagmentation regime, confirming that the protein induces additional DNA conformations, as expected. Our approach to fitting *J*-looping curves as the sum of WLC and local generic Lorentzian tagmentation functions does not allow confident analysis of very short DNA loops. We also note that *J*-loop values for the largest DNA loops in our experiments are subject to artifacts that we attribute to Tnp steric interactions with the streptavidin-modified magnetic bead surface present in the assay.

The current tagmentation-based approach thus has both obvious advantages and some limitations. It is important to emphasize that anomalous hypertagmentation for short DNA loops in the range of ∼100 bp *is no greater an obstacle than that faced for T4 DNA ligase cyclization experiments, where cyclization in this DNA length range is too slow to be quantified*. In this sense, LOOP-TAG suffers from a blind spot very comparable to existing methods, yet can determine thousands of *J*-factors at once, as well as determining the DNA persistence length in the absence or presence of additional accessory proteins of interest. Moreover, LOOP-TAG measures biologically-relevant protein-mediated DNA looping, whereas DNA cyclization experiments are arguably more artificial. It is possible to imagine LOOP-TAG variations where the DNA target terminus carries a very small reactive species, e.g. a single reactive primary amine residue such that DNA loop sites captured by formaldehyde crosslinking are detected as stops by extension of a primer hybridized near the bead-linked terminus of the target DNA.

When we focus our Worm-like Chain model fitting to the majority of the length-dependent data we find that measured *J*-loop values for annealed Tnp conjugates and antibody-mediated Tnp conjugates predict protein radii of 12.5 nm and 17.5 nm, respectively, based on our prior modeling regime (1). Interestingly, we had independently estimated from approximate molecular modeling that these radii might be ∼7 nm and ∼21 nm, respectively.

Though details of the described LOOP-TAG design can undoubtedly be optimized, we deliberately limited ourselves to commercially-available components (beads, DNA constructs, linkers, Tn5 Tnp formulations) and relatively inexpensive nanopore sequencing methodology to encourage others to explore the method. In particular, LOOP-TAG can be applied to any long DNA sequences (e.g. 1 kbp and longer) from any source, and can be used to explore the properties of relevant sequence-specific or nonspecific proteins of interest on the looping behavior of the sequence. Furthermore, there is the potential to change the looping protein itself through other DNA conjugation or annealing strategies.

## Acknowledgments

This work was supported by the Mayo Foundation and by NIH grant R35GM143949 to L.J.M. The authors acknowledge the important technical contributions of Lauren Mogil, David Tse, and Anders Herfindahl-Quint to early phases of this work, and the encouragement of Tamas Ordog to exploit tagmentation. Composition of some figures was facilitated by BioRender.com. It was the inspiring teaching of M.T. Record, Jr. more than 40 years ago that led L.J.M. to study DNA flexibility.

## Data sharing

Essential data are provided in the main manuscript and *Supplemental*. Constructs are available from the authors upon request

