## Supplemental information for "LOOP-TAG: massively-parallel measurement of length-dependent protein-mediated DNA looping probabilities by tagmentation *in vitro*"

**Table of contents**

Table S1. Buffers.

Table S2. Target DNA sequences.

Table S3. Plasmids and primers for DNA target synthesis.

Table S4. DNA target synthesis primer sequences.

Table S5. Plasmid-primer combinations for production of target DNA and PCR amplification of tagmentation products.

Table S6. Primer sequences for production of target DNA and PCR amplification of tagmentation products.

Table S7. Adapter strands for Tnp conjugation.

Table S8. qPCR primers.

Table S9. qPCR primer sequences.

Table S10. Summary of parameters from data and fitting.

Fig. S1: Overview of modeling approach.

Fig. S2: Evidence for efficient adapter loading into Tnp protein constructs.

Fig. S3: Validation of proportionality between Tnp exposure, tagmentation product mass yield, and nanopore sequencing counts.

Fig. S4. Effect of Nhp6A on Tnp activity and specificity.

### Supplemental Tables

**Table S1.** Buffers

| **Buffer** | **Composition** |
|---|---|
| B&W buffer | 5 mM Tris-HCl pH 8.0, 1 M NaCl, 0.5 mM EDTA |
| Dig X [NaCl] buffer | 20 mM HEPES pH 7.5, X mM NaCl, 0.5 mM spermidine, 0.01% Digitonin-DMSO, 1 mM EDTA pH 8.0 |
| 5× adapter loading buffer | 250 mM HEPES pH 7.5, 500 mM NaCl, 5 mM EDTA |
| 1× TE, 0.1% Triton-X buffer | 10 mM Tris-HCl pH 8.0, 1 mM EDTA, 0.1% Triton-X |
| Antibody buffer | 20 mM HEPES pH 7.5, 150 mM NaCl, 0.5 mM spermidine, 2 mM EDTA, 100 $\mu$g/mL BSA |
| Looping Buffer | 50 mM Tris-HCl pH 7.5, 1 mM EDTA |
| 10× annealing buffer | 400 mM Tris-HCl pH 8.0, 500 mM NaCl, 10 mM EDTA |
| Tnp dilution buffer | 50 mM Tris-HCl pH 7.5, 1 mM EDTA 0.1% Triton-X, 1 mM DTT, 100 mM NaCl, 50% glycerol |

**Table S2**. Biotinylated template target DNA sequences (top strand of duplex).

| Target | Plasmid source | Length (bp) | Sequence (5'-3') |
|---|---|---|---|
| 1 | pJ1506 | 1110 | **B**GCTATCAAGCTTGCTCGAGTCTCGTGGGCTCGGGCTCTGCTAATCCTGTTACCAGTGGCTGCTGCCAGTGGCGATAAGTCGT<br>GTCTTACCGGGTTGGACTCAAGACGATAGTTACCGGATAAGGCGCAGCGGTCGGGCTGAACGGGGGGTTCGTGCACACAGCCC<br>AGCTTGGAGCGAACGACCTACACCGAACTGAGATACCTACAGCGTGAGCTATGAGAAAGCGCCACGCTTCCCGAAGGGAGAAA<br>GGCGGACAGGTATCCGGTAAGCGGCAGGGTCGGAACAGGAGAGCGCACGAGGGAGCTTCCAGGGGGAAACGCCTGGTATCTTT<br>ATAGTCCTGTCGGGTTTCGCCACCTCTGACTTGAGCGTCGATTTTTGTGATGCTCGTCAGGGGGGCGGAGCCTATGGAAAAAC<br>GCCAGCAACGCGGCCTTTTTACGGTTCCTGGCCTTTTGCTGGCCTTTTGCTCACATGTTCTTTCCTGCGTTATCCCCTGATTC<br>TGTGGATAACCGTATTACCGCCTTTGAGTGAGCTGATACCGCTCGCCGCAGCCGAACGACCGAGCGCAGCGAGTCAGTGAGCG<br>AGGAAGCGGAAGAGCGCCCAATACGCAAACCGCCTCTCCCCGCGCGTTGGCCGATTCATTAATGCAGCTGGCACGACAGGTTT<br>CCCGACTGGAAAGCGGGCAGTGAGCGCAACGCAATTAATGTGAGTTAGCTCACTCATTAGGCACCCCAGGCTTTACACTTTAT<br>GCTTCCGGCTCGTATGTTGTGTGGAATTGTGAGCGGATAACAATTTCACACAGGAAACAGCTATGACCATGGCGGCGCCCGGC<br>TTCATCTCATCTGCGATGCGATAGCGTAGCGTCAGTCCAGTCTGCAGTGCAGCTGGTCTGGTAGCATAGCATCACGTCACGTG<br>TCCTGTCCTGAGGTGAGGTTGCATTGCATACAGTACAGTCAGCTCAGCTAGCGTAGCGTCTGATCTGATCGACTCGACTAACG<br>TAACGTGTCATGTCATAGGCTAGGCTCGAGTCGCGGGCGCCGCGAGAGGATCCCCGGGTACCGAGCTCGAATTCACTGGCCGT<br>CGTTTTACAAC***GTCGTGACTGGGAAAACCCTG*** |
| 2 | pJ3054 | 1161 | **B**GCTATCAAGCTTGCTCGAGTCTCGTGGGCTCGGCAGGCTTTACACTTTATGCTTCCGGCTCGTATGTTGTGTGGAATTGTGA<br>GCGGATAACAATTTCACACAGGAAACAGCTATGACCATGATTACGCCAAGCTTCGAGTCGAGTCACGTCACGTACGCAACGCA<br>TTACGTTACGCGATGCGATGTGCGTTGCGTACGCTACGCTTGTCGTGTCGCGAAGCGAAGTCCGTTCCGTACGATACGATATC<br>CGATCCGCGATTCGATTAGCGTAGCGTGCGTAGCGTAACTCGACTCGCGTACCGTACACCGTACCGTACGAAACGAAGAACGG<br>AACGCGTTGCGTTGATCGTATCGTGCGATGCGATAGTCGAGTCGCGATCCGATCAAGCGAAGCGCGTCACGTCAAACGAAACG<br>AACGACACGACGATCGGATCGCGTTTCGTTTTGCGATGCGATCGGTTCGGTATACGATACGCGTGTCGTGTTTCGTTTCGTTC<br>GAGTCGAGAAACGAAACGCGCATCGCATCTCGACTCGATCGATTCGATAACGGAACGGTCGACTCGACTTTCGTTTCGCGTGA<br>CGTGAATCGCATCGCTCGTATCGTATGACGTGACGCGTTCCGTTCATCGGATCGGTCGGATCGGATACCGTACCGCGTTACGT<br>TATACGATACGAGCGAAGCGAACTTCGCTTCGCGGAACGGAATACGTTACGTACGTCACGTCGTTCGGTTCGCGGTACGGTAA<br>CCGAACCGACCGAACCGAAATGCGATGCGCGCTACGCTAAGCGAAGCGACCGATCCGATTCACGTCACGCGAACCGAACCACG<br>ACACGACCGTACCGTAACACGACACGCGACTCGACTCTCGTCTCGTCCGTTCCGTTTATCGTATCGCGTAACGTAAATCGAAT<br>CGAACGGAACGGATTGCGTTGCGCGTAGCGTAGTACGGTACGGACGTAACGTAATTCGATTCGCGTATCGTATAACGCAACGC<br>TCGCTTCGCTAATCGAATCGCGAAACGAAATACGCTACGCTCGTTTCGTTCATCGCATCGCGGATCGGATTTCGATTCGATCG<br>TCTCGTCAACCGAACCGCGCTTCGCTTTCCGATCCGAACGAGACGAGAGACGAGACG***GAATTCACTGGCCGTCGTTTTACAAC*** |
| 3 | pJ2656 | 1033 | **B**GCTATCAAGCTTGCTCGACCTACCTGCCTTTGCGCTGGGTCGGTTACGGCCAGGACAGTCGTTTGCCGTCTGAATTTGACCT<br>GAGCGCATTTTTACGCGCCGGAGAAAACCGCCTCGCGGTGATGGTGCTGCGTTGGAGTGACGGCAGTTATCTGGAAGATCAGG<br>ATATGTGGCGGATGAGCGGCATTTTCCGTGACGTCTCGTTGCTGCATAAACCGACTACACAAATCAGCGATTTCCATGTTGCC<br>ACTCGCTTTAATGATGATTTCAGCCGCGCTGTACTGGAGGCTGAAGTTCAGATGTGCGGCGAGTTGCGTGACTACCTACGGGT<br>AACAGTTTCTTTATGGCAGGGTGAAACGCAGGTCGCCAGCGGCACCGCGCCTTTCGGCGGTGAAATTATCGATGAGCGTGGTG<br>GTTATGCCGATCGCGTCACACTACGTCTGAACGTCGAAAACCCGAAACTGTGGAGCGCCGAAATCCCGAATCTCTATCGTGCG<br>GTGGTTGAACTGCACACCGCCGACGGCACGCTGATTGAAGCAGAAGCCTGCGATGTCGGTTTCCGCGAGGTGCGGATTGAAAA<br>TGGTCTGCTGCTGCTGAACGGCAAGCCGTTGCTGATTCGAGGCGTTAACCGTCACGAGCATCATCCTCTGCATGGTCAGGTCA<br>TGGATGAGCAGACGATGGTGCAGGATATCCTGCTGATGAAGCAGAACAACTTTAACGCCGTGCGCTGTTCGCATTATCCGAAC<br>CATCCGCTGTGGTACACGCTGTGCGACCGCTACGGCCTGTATGTGGTGGATGAAGCCAATATTGAAACCCACGGCATGGTGCC<br>AATGAATCGTCTGACCGATGATCCGCGCTGGCTACCGGCGATGAGCGAACGCGTAACGCGAATGGTGCAGCGCGATCGTAATC<br>ACCCGAGTGTGATCATCTGGTCGCTGGGGAATGAATCAGGCCACGGCGCTAATCACGACGCGCTGTATCGCTGGATCAAATCT<br>GTCGATCCTTCCCGCCCG***GTGCAGTATGAAGGCGGCGG*** |

"**B**": 5' TEG-linked biotin
Underline: LOOP-TAG full-length forward primer sequence.
**Bold**: LOOP-TAG complement full-length reverse primer sequence
Target 3: non-targeted bystander DNA for studies verifying sparse magnetic bead loading

**Table S3.** Plasmids and primers for DNA target synthesis

| target | plasmid template (PCR primers) | distal terminus | forward primer | reverse primer |
|---|---|---|---|---|
| 1 | 1506 (7061/7906) | anneal | B-7061 | 7906 |
| | 1506 (7061/7063) | dig | B-7061 | 7063 |
| | 1506 (7061/7064) | blunt | B-7061 | 7064 |
| 2 | 3054 (7677/7907) | anneal | B-7677 | 7907 |
| | 3054 (7677/7612) | dig | B-7677 | 7612 |
| | 3054 (7677/7611) | blunt | B-7677 | 7611 |
| 3 | 2656 (7136/7134) | blunt | B-7136 | 7134 |

B: biotin

**Table S4**. DNA target synthesis primer sequences

| Primer | Sequence (5'-3') |
|---|---|
| 7061 | /5BiotinTEG/GCTATCAAGCTTGCTCGAGTCTCGTGGGCTCGGGCTCTGCTAATCCTGTTACCAGTG |
| 7063 | /5DigN/CAGGGTTTTCCCAGTCACGAC |
| 7064 | CAGGGTTTTCCCAGTCACGAC |
| 7134 | CCGCCGCCTTCATACTGCAC |
| 7136 | /5BiotinTEG/GCTATCAAGCTTGCTCGACCTACCTGCCTTTGCGCTGGGTCGGTTACGGCCAG |
| 7611 | GTTGTAAAACGACGGCCAGTGAATTC |
| 7612 | /5DigN/GTTGTAAAACGACGGCCAGTGAATTC |
| 7613 | /5DigN/CCGCCGCCTTCATACTGCAC |
| 7677 | /5BiotinTEG/GCTATCAAGCTTGCTCGAGTCTCGTGGGCTCGGCAGGCTTTACACTTTATGCTTCCGGCTC |
| 7906 | ctgccgacga/iSp9/CAGGGTTTTCCCAGTCACGAC |
| 7907 | ctgccgacga/iSp9/GTTGTAAAACGACGGCCAGTGAATTC |

Lower case: remains single-stranded after PCR
/5BiotinTEG/: IDT biotin with tri(ethylene glycol) spacer
/5DigN/: IDT digoxigenin coupled with NHS ester chemistry
/iSp9/: IDT non-extendable spacer

**Table S5**. Plasmid-primer combinations for production of target DNA and PCR amplification of tagmentation products

| **Plasmid** | **Forward Primer** | **pR (Full Length)** | **pR (Tagmentation)** |
|---|---|---|---|
| 1506 (target 1) | 7873 | 7064 | 7828 |
| 3054 (target 2) | 7877 | 7611 | 7828 |
| 2656 (target 3) | 7876 | 7134 | 7828 |

**pR:** reverse primer

**Table S6**. Primer sequences for production of target DNA and PCR amplification of tagmentation products

| Primer | Sequence (5'-3') |
|---|---|
| 7064 | CAGGGTTTTCCCAGTCACGAC |
| 7134 | CCGCCGCCTTCATACTGCAC |
| 7611 | GTTGTAAAACGACGGCCAGTGAATTC |
| 7828 | GTCATCGTCGGCAGCGTC |
| 7873 | GGCTCGGGCTCTGCTAATC |
| 7876 | ACCTGCCTTTGCGCTGGGTC |
| 7877 | GGCAGGCTTTACACTTTATGCTTC |

**Table S7**. Mosaic A adapter strands for Tnp loading

| Strand | Sequence (5'-3') |
|---|---|
| 7065 | TCGTCGGCAGCGTCAGATGTGTATAAGAGACAG |
| 7066 | /5Phos/CTGTCTCTTATACACATCT |
| 7901 | /5Phos/CTGTCTCTTATACACATCTGACG |
| 7987 | CGTTTCCGTCCATCCCCGAGCCCACGAGACAGATGTGTATAAGAGACAG |

/5Phos/: IDT phosphorylation

**Table S8**. qPCR primers

| Template | Forward Primer | Reverse Primer |
|---|---|---|
| 1506 | 7873 | 7676 |
| 3054 | 7877 | 7674 |
| 7897/7066 Adapter | 7908 | 7900 |

**Table S9**. qPCR primer sequences

| Primer | Sequence (5'-3') |
|---|---|
| 7674 | CGAAGCTTGGCGTAATCATG |
| 7676 | ACCGCTGCGCCTTATCCG |
| 7873 | GGCTCGGGCTCTGCTAATC |
| 7877 | GGCAGGCTTTACACTTTATGCTTC |
| 7900 | ACGTTTCCGTCCATC |
| 7908 | GCAGCTGTCTCTTATACACATCTG |

**Table S10.** Summary of parameters from data and fitting

| Parameter | Annealed Tnp | Antibody-conjugated Tnp |
|---|---|---|
| Fraction of assembled Tnp: *f* | 0.0085 – 0.0315 | 0.0079 – 0.0201 |
| Minimum *J*-loop | $9.1 \times 10^{-9}$ M | $1.7 \times 10^{-8}$ M |
| Loop length at minimum *J* -loop | 47 bp | 137 bp |
| Maximum *J* -loop | $1.7 \times 10^{-7}$ M | $5.0 \times 10^{-6}$ M |
| Loop length at maximum *J* -loop | 23 bp | 2 bp |
| Maximum *J* -loop with Nhp6A | $2.2 \times 10^{-5}$ M | $2.0 \times 10^{-6}$ M |
| **Left-side Lorentzian model** | | |
| Peak amplitude: *A* | $1.7 \times 10^{-7}$ M | $5.0 \times 10^{-6}$ M |
| Peak center: $\mu$ | 16.4 bp | 2.9 bp |
| Half width at half max: $\gamma$ | 7.0 bp | 3.4 bp |
| **Right-side Lorentzian model** | | |
| Peak amplitude: *A* | $2.2 \times 10^{-5}$ M | $5.0 \times 10^{-6}$ M |
| Peak center: $\mu$ | 0 bp | 0 bp |
| Half width at half max: $\gamma$ | 1.3 bp | 8.6 bp |
| **Protein–mediated WLC model** | | |
| Persistence length: *P* | 50 nm | 50 nm |
| Protein bridge radius: *r* | 12.5 nm (~39 bp) | 17.4 nm (~55 bp) |
| Kink angle: $\kappa$ (180° : no kink) | 180° | 180° |
| **Summation model** | | |
| Excess *J*-loop area under Lorentzian curve | $3.8 \times 10^{-6}$ M·bp | $6.6 \times 10^{-5}$ M·bp |

### Supplemental Figures

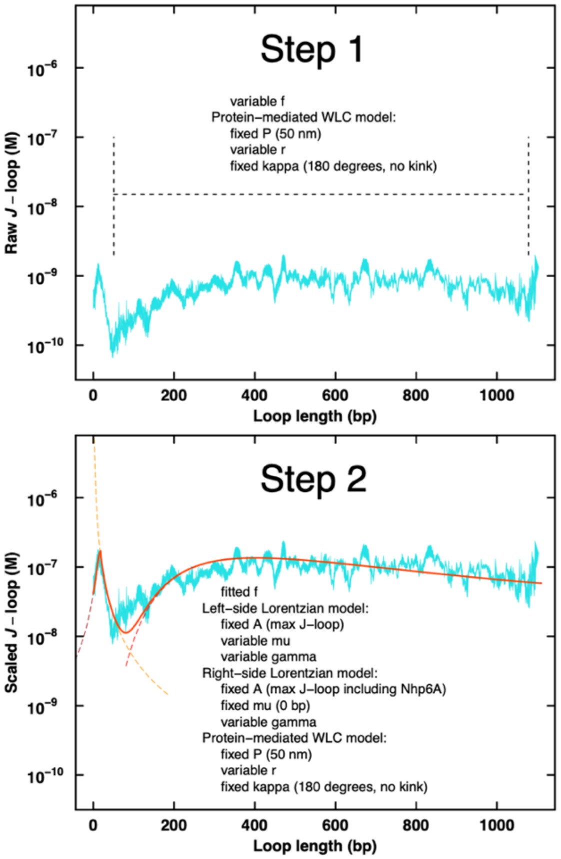


Protein-mediated WLC model

$$C_{\mathrm{SY}} = \frac{1.66}{P^3} \times \frac{112.04}{(L/P)^5} \times e^{0.246(L/P)}$$

$$J_{\mathrm{protein}} = C_{\mathrm{SY}}(L + 2r) \times e^{\left(\frac{7.1-0.1155\kappa}{(L+2r)/P}\right)}$$

$P$: persistence length
$r$: protein bridge radius
$\kappa$: kink angle

Lorentzian model

$$y = A\left(\frac{1}{1+\left(\frac{x-\mu}{\gamma}\right)^2}\right)$$

$A$: peak amplitude (maximum peak height)
$\mu$: center of the peak
$\gamma$: half width at half maximum

**Fig. S1**. Overview of modeling approach. Fitting was divided into two steps. The first step utilized only the protein-mediated Worm-like Chain (WLC) model and was designed to determine the fractional assembly parameter (*f*) for the efficiency of assembling adapter-loaded Tnp conjugates on ~1,000-bp bead-immobilized DNA. Two of the three additional parameters in this step were fixed. Additionally, the fitting was confined to loop lengths that avoided the anomalous very short-length high-probability tagmentation regime and the very long-length bead-proximal regime with higher background signal. The fitted values of the fractional assembly parameter (*f*) were then fixed for the second step. The second step involved fitting the now corrected (scaled by *f*) mean *J*-loop data over the entirety of loop lengths using a summation model to include the anomalous short-length tagmentation regime. This summation model consisted of a generic Lorentzian function divided into two sides and summed with the protein-mediated WLC model.

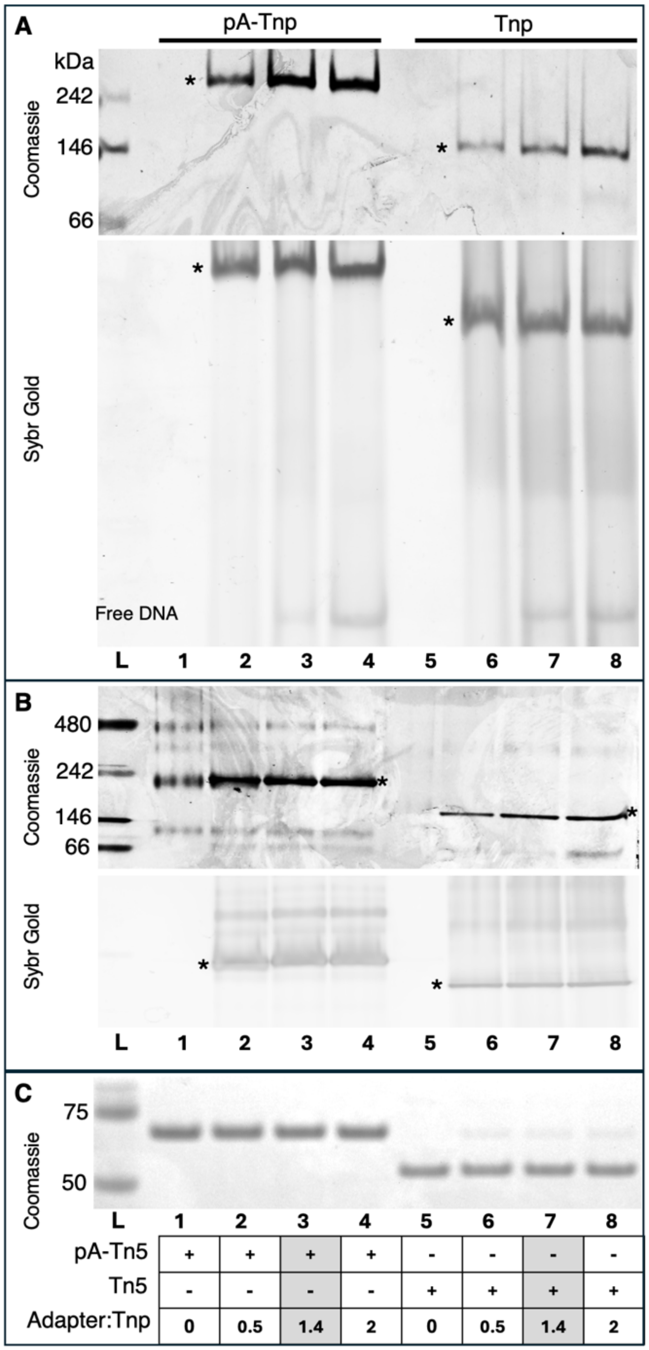


**Fig. S2.** Evidence for efficient adapter loading into Tnp protein constructs. A-C. pA-Tnp (lanes 1-4) or Tnp (lanes 5-8) were loaded with increasing molar ratios of adapter A (7065/7066) : Tnp (0:1, 0.5:1, 1.4:1, and 2:1). Identical samples were loaded onto a 6% 29:1 acrylamide:bisacrylamide native polyacrylamide gel (A), a 3-12% native blue polyacrylamide gradient gel (B) or a 10% bis-Tris SDS denaturing polyacrylamide gel (C). Gels were post-stained first with SYBR Gold, then with Coomassie protein stain (indicated at left). Adapter A-loaded Tnp dimer complexes are indicated (*) in (A) and (B). The mosaic adapter A:Tnp 1:1.4 ratio used for nanopore sequencing studies is indicated (lanes 3 and 7, gray shading). Protein MW markers (L) are shown with indicated size (kDa).

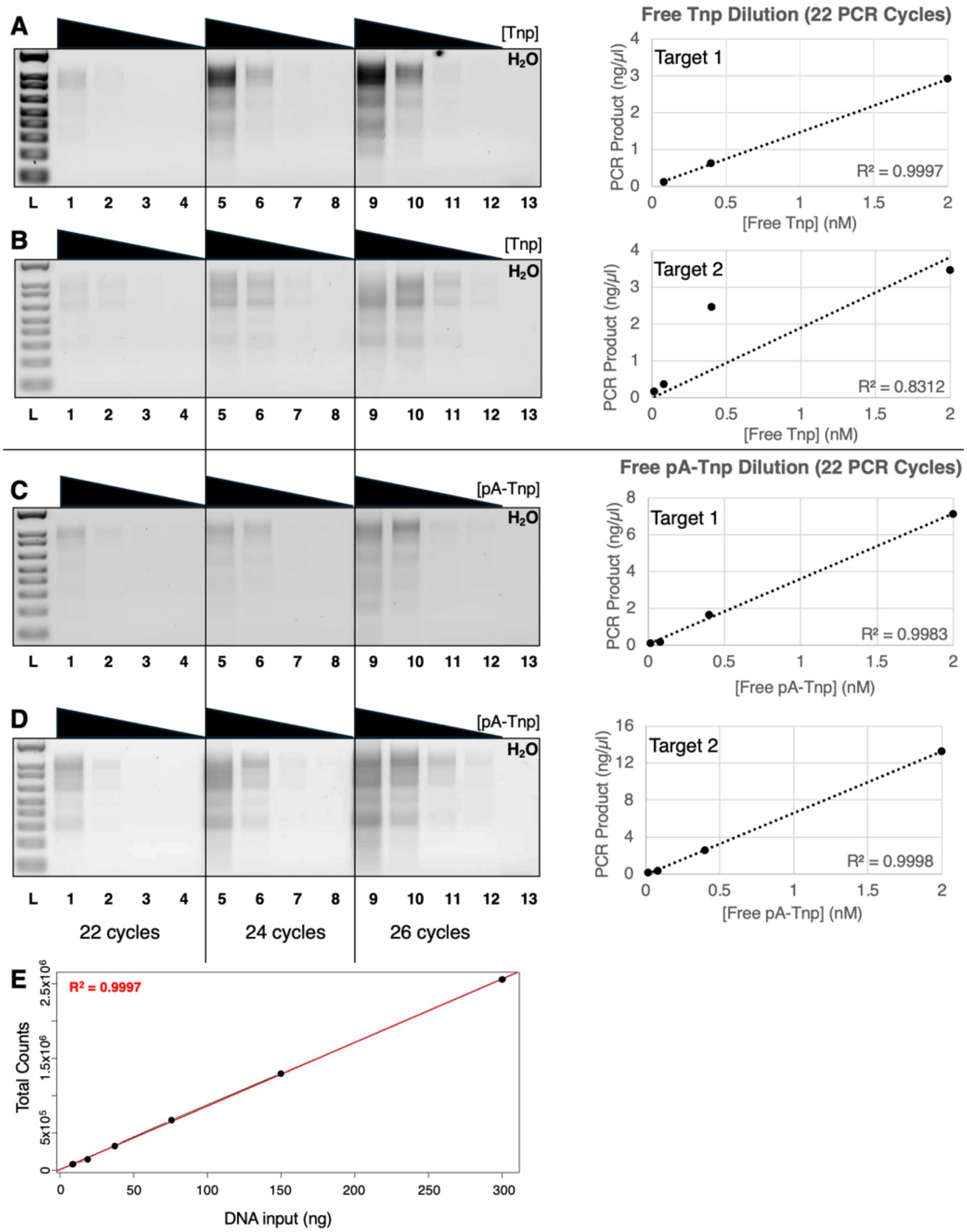


**Fig. S3**. Validation of proportionality between Tnp exposure, tagmentation product mass yield, and nanopore sequencing counts. A-D. Tagmentation by free Tnp or pA-Tnp. Targets 1 and 2 were exposed to 2 nM, 0.4 nM, 0.08 nM, or 0.016 nM Tnp or pA-Tnp followed by PCR amplification for the indicated cycle numbers (below). Purified PCR products were visualized on a 2% agarose gel (left), Qubit quantification of recovered PCR product mass is shown (right) after 22 PCR cycles. Panels A and B show results for free Tnp. Panels C and D show results for free pA-Tnp. E. Proportionality of PCR amplified and purified tagmented Target 3 DNA to nanopore sequencing counts across two-fold dilutions (ranging from 300 ng to 9.375 ng) after conversion to bar-coded nanopore sequencing libraries.

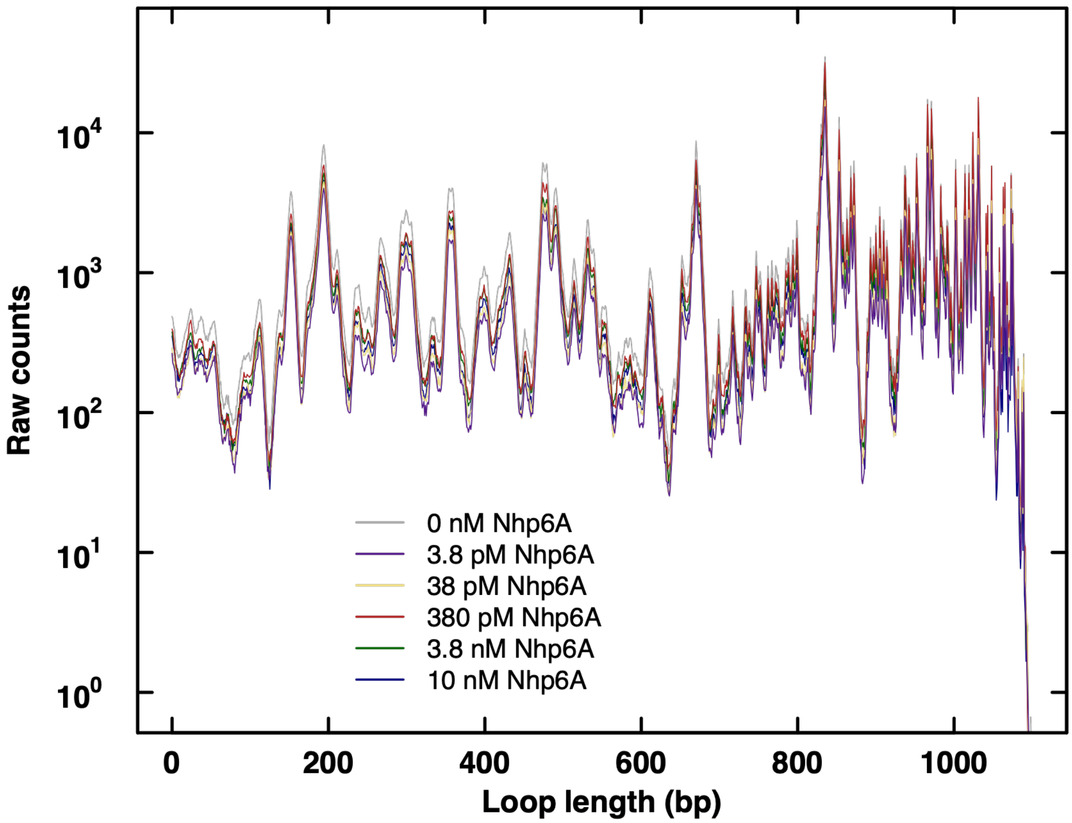


**Fig. S4**. Minimal effect of Nhp6A on Tnp activity and specificity. Free pA-Tnp tagmentation (2 nM nominal concentration) was performed on Target 2 in the presence of the indicated concentrations of Nhp6A to determine if the architectural protein would modify tagmentation activity or specificity. Because there was minimal effect of Nhp6A on Tnp activity (mean of three replicates shown), data for free Tnp in the absence of Nhp6A were used for *J*-loop calculations of Nhp6A effects on looping of tethered DNA.
